# Ketogenic Diet Enhances Cognitive-Behavioral Function and Hippocampal Neurogenesis While Attenuating Amyloid Pathology in Tg-SwDI Mice

**DOI:** 10.1101/2025.02.19.639138

**Authors:** Victoria E. Pulido-Correa, Ariana Hernandez, Eleanor J. Wind, YingYing Zhu, Chana Vogel, Shaina Binu, Mikayla Jeneske, Dylan Kuni, Vedika Chiduruppa, Bianca Echeverria, Lauren Rosenberg, Lisa S. Robison

## Abstract

The ketogenic diet (KD), characterized by high-fat, low-carbohydrate, and moderate protein intake, has gained attention for its therapeutic potential in patients with neurodegenerative diseases, including Alzheimer’s disease. Studies in Alzheimer’s rodent models report that KD and/or ketogenic supplements attenuate cognitive-behavioral impairments, neuroinflammation, amyloid-beta plaques and tau pathology. However, it is unknown whether KD can similarly benefit individuals with cerebral amyloid angiopathy (CAA), a prevalent condition in which amyloid accumulates in cerebral vessels. CAA is highly comorbid in patients with Alzheimer’s and, on its own, increases the risk of stroke, cognitive impairment, and dementia, yet no effective treatments currently exist. The objective of this study was to determine whether KD can improve cognitive-behavioral and neuropathological outcomes in a mouse model with CAA. Male Tg-SwDI mice were fed either a standard chow or KD from 3.5 to 7.5 months of age. Following ∼3 months of dietary intervention, glucose and ketone-body levels were assessed, then mice underwent a battery of behavioral tests to evaluate locomotor activity, anxiety-related behaviors, and cognition. Immunohistochemistry was performed to assess amyloid pathology, vascular density, neuroinflammation, white matter integrity, and hippocampal neurogenesis. In addition to KD inducing nutritional ketosis and achieving metabolic benefits, mice on KD exhibited increased activity, enhanced spatial learning and memory, and a trend toward improved spatial working memory. These cognitive benefits were accompanied by an attenuation of amyloid pathology and increased hippocampal neurogenesis. These findings suggest that a ketogenic diet may be safe and effective in Alzheimer’s and dementia patients with CAA.

## INTRODUCTION

Cerebral amyloid angiopathy (CAA), the accumulation of amyloid protein (most commonly amyloid-β; Aβ) in the cerebral vasculature, is one of the most prevalent forms of cerebral small vessel disease. Most CAA cases are sporadic and associated with aging, though familial forms exist. As a single pathology, CAA can result in vascular cognitive impairment and dementia (VCID) and significantly increases the risk of stroke.1–3 One of the most significant clinical consequences of CAA is the increased risk of intracerebral hemorrhages. These can be large or small (microbleeds), and recurrent hemorrhages are common. While hemorrhages are more common, CAA can also lead to ischemic events, including transient ischemic attacks and strokes. The progressive accumulation of amyloid in vessel walls can narrow the lumen (interior space) of the vessels, leading to chronic hypoperfusion. This reduced blood flow can cause gradual damage to brain tissue, contributing to cognitive decline. Indeed, CAA is specifically linked to impairments in several cognitive domains, including perceptual speed and episodic memory.1 CAA can also be associated with inflammation characterized by activated microglia and reactive astrocytes that produce pro-inflammatory cytokines and chemokines, reactive oxygen and nitrogen species, and activation of the complement pathway.3 Of note, CAA is commonly found in patients with Alzheimer’s disease;4 this overlap can exacerbate cognitive decline and other neurological symptoms.5 The high rate of comorbidity is perhaps unsurprising, as both CAA and Alzheimer’s disease are characterized by the accumulation of Aβ, albeit differing in their distribution (blood vessel walls in CAA vs. brain parenchyma in Alzheimer’s). The processes that govern Aβ clearance and deposition are common to both diseases. Indeed, CAA likely contributes to Alzheimer’s pathology by impairing perivascular Aβ drainage from the brain, a major route of Aβ clearance.6, 7

Currently, there is no cure for CAA, nor are there any FDA-approved treatments. Recently approved Aβ-directed monoclonal antibodies (e.g. lecanemab) facilitate parenchymal Aβ removal in patients with Alzheimer’s; however, these treatments have questionable benefits for cognition and carry substantial risk for amyloid-related imaging abnormalities (ARIA), including brain swelling (edema) and bleeding (hemorrhage).8,9–12 ARIA resulting from the treatment of Alzheimer’s with Aβ-directed monoclonal antibodies may be of particular concern in individuals with CAA.9–13 In fact, it was recently hypothesized that patients positive for CAA should be identified and excluded from anti-amyloid therapy (AAIC Proceedings). Therefore, novel interventions are necessary for patients with CAA, either as a singular pathology or in cases where CAA is comorbid with Alzheimer’s disease.

One potential nonpharmacological intervention for CAA that is yet to be tested is the ketogenic diet, which involves the consumption of limited carbohydrates (usually <5%), moderate protein intake, and very high fat.14 Limiting the intake of carbohydrates induces a state of “nutritional ketosis” that promotes ketogenesis while reducing gluconeogenesis.14 This causes ketone bodies to replace glucose as the body’s primary energy source. Originally used for the treatment of epilepsy, the ketogenic diet has gained considerable attention in recent decades for its ability to produce quick weight loss and improve metabolic outcomes, such as those related to prediabetes/Type II diabetes (e.g., insulin resistance, blood glucose).15, 16 Additionally, growing evidence supports the efficacy of ketogenic diet to promote brain health and improve cognitive performance across a number of domains, such as working memory, reference memory, and attention.17 Notably, the ketogenic diet is found to be protective in a variety of neurodegenerative diseases, including Alzheimer’s disease, Parkinson’s disease, and motor neuron disease.18 Emerging research suggests that consumption of a ketogenic diet or supplementation with exogenous ketones exerts many beneficial effects, including cognitive-boosting, pro-neurogenic, anti-inflammatory, and antioxidant effects.19–22 Studies in Alzheimer’s rodent models have found that ketogenic diet and/or exogenous treatment with ketone supplements attenuate cognitive-behavioral impairments, neuroinflammation, oxidative stress, and Aβ accumulation; and enhance mitochondrial function and synaptic plasticity.21 A review of the use of either ketogenic diet or exogenous ketone supplementation reported that these ketogenic dietary interventions provide treatment benefits in Alzheimer’s patients; however, most studies are small and uncontrolled with relatively short-term treatment periods.23

There is evidence to suggest that ketogenic dietary interventions promote brain health and boost cognitive performance, are protective against neurodegenerative diseases, including Alzheimer’s disease, and aid in the recovery from cerebrovascular events, such as stroke.17, 19, 21, 23–25 However, it is yet to be determined whether ketogenic diet can similarly benefit patients with CAA. Here, we used a transgenic mouse model with CAA (Tg-SwDI mice) to assess the ability of the ketogenic diet to improve cognitive-behavioral and neuropathological outcomes during early-to-moderate stage disease.

## METHODS

### Animals and housing conditions

All experiments were approved by the Institutional Animal Care and Use Committee at Nova Southeastern University (Davie, FL, USA) and compliance was maintained with the ARRIVE guidelines.26 Male Tg-SwDI mice (JAX MMRRC stock #034843) were bred for this experiment at Nova Southeastern University using mice purchased from Jackson Laboratory (Bar Harbor, Maine). These mice carry three mutations in the human amyloid-β protein precursor (AβPP) gene: Iowa (APP/D694N), Dutch (APP/E693Q), and Swedish (K670N/M671L). These mutations lead to the accumulation of fibrillar amyloid beta peptides in the cerebral vasculature, effectively replicating the key pathological features of CAA.27 Tg-SwDI mice also exhibit cerebrovascular dysfunction, neuroinflammation, and cognitive-behavioral impairment, as we and others have shown previously.27–32 Animals were housed in groups of three to four in standard polysulfone cages (dimensions: 7.75"L x 14.75"W x 5.25"H). The housing facility was kept on a reversed 12- hour light-dark cycle (lights on at 8:00 PM and off at 8:00 AM) to mimic the natural circadian rhythm of the mice. Temperature and humidity were set at 68–72 °F and 40-60%, respectively. These conditions were monitored and recorded daily to ensure a stable and comfortable environment conducive to the mice’s physiological needs. Mice were given ad libitum access to fresh food and water and were provided with various environmental enrichment items (plastic huts, chewing bones, nesting materials) for the duration of the study. All work with mice was performed during the dark cycle.

### Diet intervention

A timeline of the experiment is shown in **Figure 1**. All mice were fed a standard chow diet until ∼3.5 months of age. At that point, mice were either switched to a ketogenic diet (n=8) or maintained on the standard chow diet (n=7) for the duration of the study (∼4 months). The Ketogenic Diet AIN-76A Modified (Catalog #F3666, Bio-Serv, Flemington, New Jersey) was composed of 93.4% fat, 4.7% protein, and 1.8% carbohydrates, and had a caloric value of 7.24 kcal/g. The standard chow diet (5P76, LabDiet, St. Louis, Missouri) was composed of 14.4% fat, 26.1% protein, and 59.5% carbohydrates. Mice were weighed and food replenished twice weekly. Average food and fluid intake per mouse were calculated for each cage based on the amount consumed and the number of mice housed in the cage; food (kcal) and fluid (mL) consumption were then normalized to body weight.

**Figure 1.**
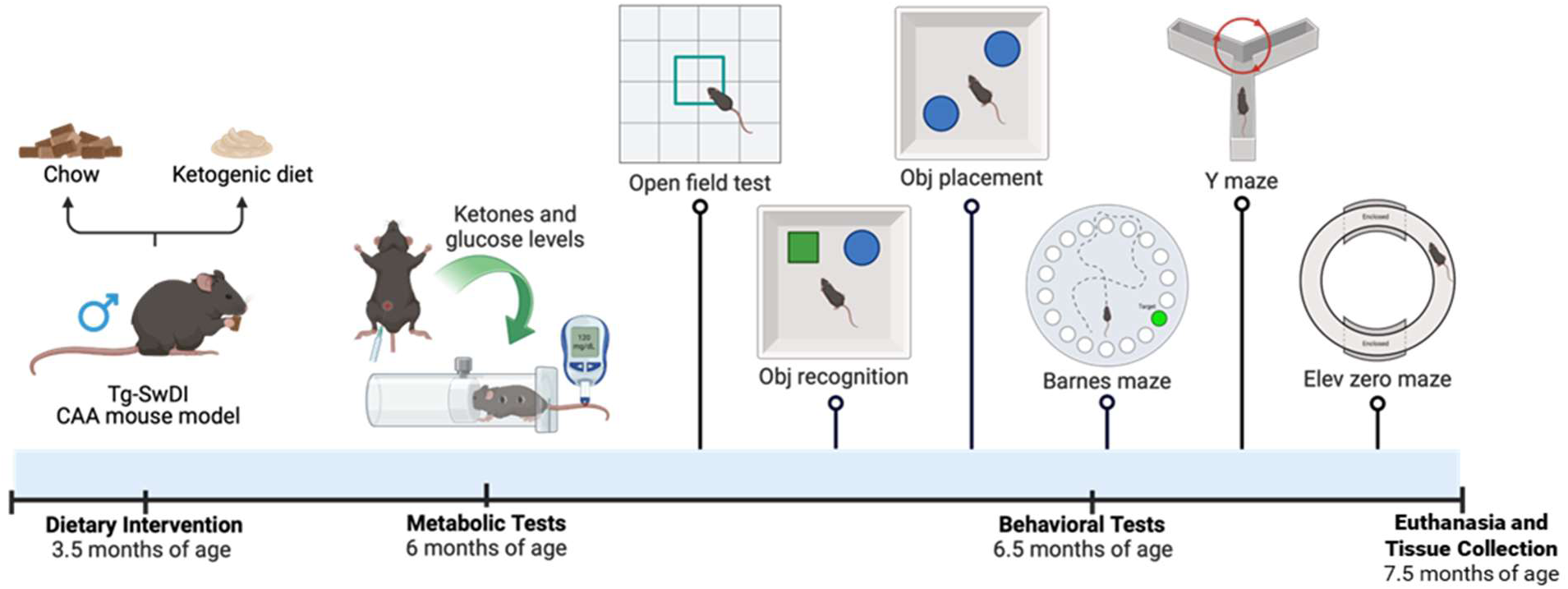
Timeline of the experiment. Male Tg-SwDI mice either remained on standard rodent chow or were switched to a ketogenic diet at ∼3.5 months of age, with diet maintained through the rest of the study. At 6 months of age, metabolic testing was performed to assess blood ketone body and glucose levels during fed and fasting state, as well as glucose tolerance. Behavior testing was performed to assess locomotor activity, anxiety-like behavior, and cognition, followed by euthanasia and tissue collection at 7.5 months of age. Figure created with Biorender.

### Ketone body levels, blood glucose levels, and glucose tolerance testing (GTT)

Ketone body levels, fasting blood glucose levels, fed blood glucose levels, and glucose tolerance were assessed starting at ∼6 months of age. Ketone body levels and fed blood glucose levels were assessed in the fed state ∼2-4 hours into the dark cycle, measured with a blood ketone meter and glucometer, respectively. Blood was collected via tail snip (<1 mm) following application of topical analgesic (Aspercreme).

Mice were given a glucose tolerance test (GTT) to assess diabetic status on a subsequent day. Mice were fasted for ∼16 hours, and their fasting blood glucose levels were measured (t = 0) using a glucometer with blood collected via tail snip (<1 mm) following application of topical analgesic (Aspercreme). Following an i.p. injection of 2 g/kg of glucose, blood glucose levels were measured again at 15, 30, 60, and 120 min post-injection to assess glucose tolerance.

### Behavior testing

Three months after diet initiation, mice were subjected to a battery of behavioral tests, including open field, novel object recognition test, object placement test, Y-maze, elevated zero maze, and Barnes maze. Mice were acclimated to the behavior testing room for at least 30 minutes prior to the start of testing. Following each trial, behavior equipment and any objects used for the test were cleaned with 70% isopropyl alcohol and allowed to dry thoroughly prior to testing.

#### Open field

Open field testing was performed to assess general activity levels/exploratory behavior and anxiety-like behavior. Mice were placed into an open field arena (46 cm x 46 cm x 46 cm; opaque white acrylic walls and floor), and behavior was recorded for 10 minutes. Distance traveled, time spent moving, speed, and center activity were measured using automated software (Noldus Ethovision v17, Wageningen, the Netherlands), while measures of rearing and grooming behavior were manually recorded by two independent raters blinded to the experimental condition.

#### Novel object recognition test (NORT)

NORT was performed to assess object recognition memory using two trials, both of which were performed in the open field arena. In the first trial (training), mice were placed in the arena with two of the same objects and allowed to explore for 10 minutes. Mice were returned to their home cage for a 1-hour retention period, after which mice were placed back into the arena for the second trial (test). In the test trial, there were two objects in the arena in the same location as the training trial. This included one “familiar” object (previously exposed to in the training trial) and one “novel” object. The time spent with each object was recorded for 5 minutes, measured using automated software (Noldus Ethovision v17, Wageningen, the Netherlands), with intact memory determined by more interaction with the novel object versus the familiar object. The discrimination index was calculated as [(time with novel object – time with familiar object)/time with both objects].

#### Object placement test (OPT)

OPT was performed to assess spatial memory using two trials, both of which were performed in the open field arena. In the first trial (training), mice were placed in the arena with two of the same objects and allowed to explore for 10 minutes. Mice were returned to their home cage for a 1-hour retention period, after which mice were placed back into the arena for the second trial (test). In the test trial, the same two objects were placed in the arena as in the training trial. This included one object in a “familiar” location (same as in the training trial) and one “displaced” object (location switched from the previous training trial). The time spent with each object was recorded for 5 minutes, measured using automated software (Noldus Ethovision v17, Wageningen, the Netherlands), with intact memory determined by more interaction with the displaced object versus the familiar object. The discrimination index was calculated as [(time with displaced object – time with familiar object)/time with both objects].

#### Y-maze

The Y-maze was performed to assess spatial working memory. Each mouse received one 5-minute trial performed in a Y-shaped maze consisting of three opaque arms at a 120° angle from each other (each arm = 30 cm L x 6 cm W x 15 cm H). The order of arm entries was recorded by an experimenter blinded to experimental conditions. The number of arm entries was counted as a measure of exploratory behavior. Rodents typically prefer to explore a new arm of the maze instead of returning to one that they have previously visited.33 Over the course of multiple arm entries, the subject should alternate the arms visited, showing a tendency to enter a less recently visited arm. The number of triads (sequence of each of the three arms visited) was recorded in order to calculate the percentage of alternation. An entry into an arm was only counted when all four paws were within that arm. Spontaneous alternation (indicative of intact spatial working memory) was calculated as ([# triads/(total # arm entries – 2)]*100).

#### Elevated zero maze

The elevated zero maze was performed to assess anxiety-like behavior. The elevated zero maze is composed of a circular “O”-shaped platform that consists of 2 closed sections (high walls surrounding the edges) and 2 open sections (no surrounding walls). Each mouse was given a single 5-minute trial that began by placing the mouse into a closed portion of the maze. Number of entries and time spent in and near the open sections were recorded as measures of reduced anxiety-like behavior measured using automated software (Noldus Ethovision v17, Wageningen, the Netherlands).

#### Barnes maze

The Barnes maze was performed to assess spatial learning and memory. The Barnes maze is an elevated circular white platform (diameter = 122 cm) with 40 evenly-spaced holes around the perimeter. One hole was deemed the “target hole,” which had a cup mounted underneath it, while the remaining 39 holes were left empty. Because bright light and open spaces are aversive, mice are motivated to locate and enter the cup under the escape hole. A piece of clean tissue paper (Kimwipe) was also placed inside the cup. The Barnes maze was surrounded by white curtains, which had visual cues attached to each. The location of the target hole, curtains, and visual cues were maintained for the duration of habituation, training, and probe trials.

Day 1 consisted of a single habituation trial. During habituation, mice were placed in the escape cup for one minute, covered by a clear glass beaker. They were then placed in the center of the arena, covered by a wrapped beaker for 15 seconds prior to the start of a 5-minute exploration period. The exploratory period lasted 5 minutes, regardless of whether the mouse entered the escape cup. At the end of the 5 minutes, mice were placed in the escape cup covered with a clear glass beaker for 1 minute, then returned to their home cage.

Days 2-4 consisted of 2 training trials per day, with daily trials separated by approximately 1 hour. Mice were placed in the center of the arena, covered by a wrapped beaker for 15 seconds prior to the start of each training trial. Mice had a maximum of 3 minutes to find and enter the escape cup, at which time the trial ended, and were covered with a clear glass beaker for 1 minute. If mice did not enter the escape cup within 3 minutes, they were gently guided to the escape cup and covered with a clear glass beaker for 1 minute, then returned to their home cage. The latency to find the escape cup over the 6 total training trials was calculated as a measure of spatial learning using automated software (Noldus Ethovision v17, Wageningen, the Netherlands).

On day 5, a single probe trial was conducted ∼24 hours after the last training trial (1-day probe). Mice were placed in the center of the arena, covered by a wrapped beaker for 15 seconds prior to the start of the trial. The escape cup was removed from the arena for the probe trials. The exploratory period lasted 2 minutes for each mouse. An additional probe trial was conducted 7 days after the last training trial to assess longer-term spatial memory (7-day probe). For analysis of each probe trial, automated tracking software (Noldus Ethovision v17, Wageningen, the Netherlands) was used to determine the primary latency (latency to first target hole visit) and the time spent in the target quadrant, where the escape cup was previously located.

### Euthanasia and tissue collection

Euthanasia was performed and tissue was collected after completion of all behavior testing. Mice were deeply anesthetized with isoflurane anesthesia (5% induction; ∼3-4% maintenance) and perfused with ice-cold 0.9% NaCl. Euthanasia was confirmed via decapitation, followed by dissection and recording of wet weights of visceral and subcutaneous fat. Brains were bisected, with one hemisphere undergoing fixation and the other hemisphere being flash-frozen. The fixed hemisphere was placed into 4% PFA for 48h, followed by cryopreservation using 30% sucrose, both at 4°C. The flash-frozen hemisphere was frozen in 2-methylbutane over dry ice, then stored at –80°C until use.

### Immunofluorescence

Fixed brain hemispheres were embedded in OCT and cut on a cryostat at ∼18°C into 4 sets of 40 µM sagittal free-floating sections. Free-floating immunohistochemistry was performed to assess the glial markers, ionized calcium-binding adapter molecule 1 (Iba1; microglia) and glial fibrillary acidic protein (GFAP; astrocytes), as well as white matter integrity (myelin basic protein; MBP) and neurogenesis (doublecortin; DCX, a measure of neuroblasts/immature neurons).

Sections were washed in 1x PBS with 0.01% sodium azide (3x 3 min at room temperature), permeabilized in 0.3% Triton X-100 in 1x PBS with 0.01% sodium azide (30 min at room temperature), blocked in 4% donkey serum with 0.3% Triton X-100 in 1x PBS with 0.01% sodium azide (30 min at room temperature), then incubated with primary antibodies overnight at 4°C. Primary antibodies were diluted in blocking buffer, which included rabbit anti-Iba1 (1:1000 Cat # 019-19741, Fujifilm Wako, Richmond, VA), rabbit anti-GFAP (1:100, Cat #PB9082, Boster Bio, Pleasanton, CA), guinea pig anti-DCX (1:1000, Cat #AB2253, Millipore, Burlington, MA), rabbit anti-MBP (1:200, Cat #10458-1-AP, Proteintech, Rosemont, IL). The next day, sections were washed (3x 3 min at room temperature), then incubated for 1h with secondary antibody solution at room temperature. Secondary antibodies were diluted in blocking buffer, which included donkey anti-guinea pig 488 (1:300, Cat #706-545-148, Jackson Immunoresearch, West Grove, PA), donkey anti-rabbit 594 (1:300, Cat #A21206, Invitrogen, Carlsbad, California), or donkey anti-rabbit 594 (1:300, Cat #711-585-152, Jackson Immunoresearch, West Grove, PA). Sections were washed with 1x PBS with 0.01% sodium azide (3x 3 min at room temperature) prior to being mounted on glass microscope slides with Prolong Gold Antifade mounting media with DAPI counterstain.

Fresh frozen brain hemispheres were cut on a cryostat at ∼13°C into 3 sets of 20 µM sagittal sections thaw-mounted onto charged glass microscope slides and stored at –80°C until use. Sections were thawed to room temperature, fixed with 4% paraformaldehyde in PBS (10 min at room temperature), washed in 1x PBS with 0.01% sodium azide (3x 3 min at room temperature), permeabilized in 0.1% Triton X-100 in 1x PBS with 0.01% sodium azide (30 min at room temperature), blocked in 4% donkey serum with 0.1% Triton X-100 in 1x PBS with 0.01% sodium azide (30 min at room temperature), then incubated with primary antibody overnight at 4°C. For the primary antibody solution, rabbit anti-collagen IV (1:200, Cat #PA1-28534, Invitrogen,

Carlsbad, California) was diluted in blocking buffer. The next day, sections were washed (3x 3 min at room temperature), then incubated for 1h with secondary antibody solution at room temperature. For the secondary antibody solution, donkey anti-rabbit 594 (1:300, Cat #A21206, Invitrogen, Carlsbad, California) was diluted in blocking buffer. Following three washes (3x 3 min at room temperature), sections were stained with 0.0125% Thioflavin-S in 50% PBS/50% EtOH (15 min at room temperature). Sections were washed with 50% PBS/50% EtOH (2x 3 min at room temperature) then 1x PBS with 0.01% sodium azide (3x 3 min at room temperature) prior to being mounted with Prolong Gold Antifade mounting media.

Images were taken using a 5x or 10x objective with cellSens Imaging Software on an Olympus IX73 fluorescence microscope. For analysis of thioflavin-S staining (amyloid), collagen IV (blood vessels), Iba1 (microglia), and GFAP (astrocytes), images were taken in the hippocampus (subiculum, dentate gyrus, CA1, and CA3), thalamus, and cortex. For analysis of MBP (myelin), images were taken in the hippocampus, corpus callosum, and cortex. For analysis of DCX (neurogenesis), images were taken in the dentate gyrus of the hippocampus. NIH ImageJ software was used to identify regions of interest, threshold images, and extract data on intensity and/or % area covered for each brain region for Thioflavin-S, Collagen IV, Iba-1, GFAP, and MBP. Colocalization analysis was also performed for Thioflavin-S/Collagen IV for the assessment of vascular (CAA) and non-vascular (parenchymal/plaque) amyloid accumulation. Plaque # was manually counted in the cortex of sections stained with Thioflavin-S, while DCX+ cells (neuroblasts/immature neurons) were manually counted in the dentate gyrus of the hippocampus as a measure of neurogenesis. DCX+ cells were counted by two independent raters who were blinded to the experimental condition of each mouse.

### Statistical analyses

Statistical analyses were performed using Prism v10.1.1 (GraphPad Software, San Diego, CA, USA). The Grubbs’ test, a widely accepted method for detecting outliers in small datasets, was conducted to detect statistically significant outliers (α = 0.05), followed by replacement with the next most extreme value. Unpaired two-sample t-tests were conducted for data that did not include repeated measures. Two-way repeated measures ANOVAs were performed for data that were collected and reported at multiple timepoints (body weight, fasting and fed blood ketone body levels, fasting and fed blood glucose levels, glucose tolerance test). One-sample t-tests were performed for analysis of NOR and OPT performance (discrimination index vs. hypothetical value of 0). The threshold for statistical significance was set at p < 0.05.

## RESULTS

### Metabolic outcomes

#### Caloric intake

Caloric intake for each cage was approximated based on food weights and energy density of respective diet; these values were then normalized by average body weight of the mouse in that cage to calculate food intake in kcal/g of body weight **(Figure 2a)**. An unpaired two-sample t-test found that there was no significant difference in caloric intake between groups [t(3)=0.092, p=0.9318].

**Figure 2.**
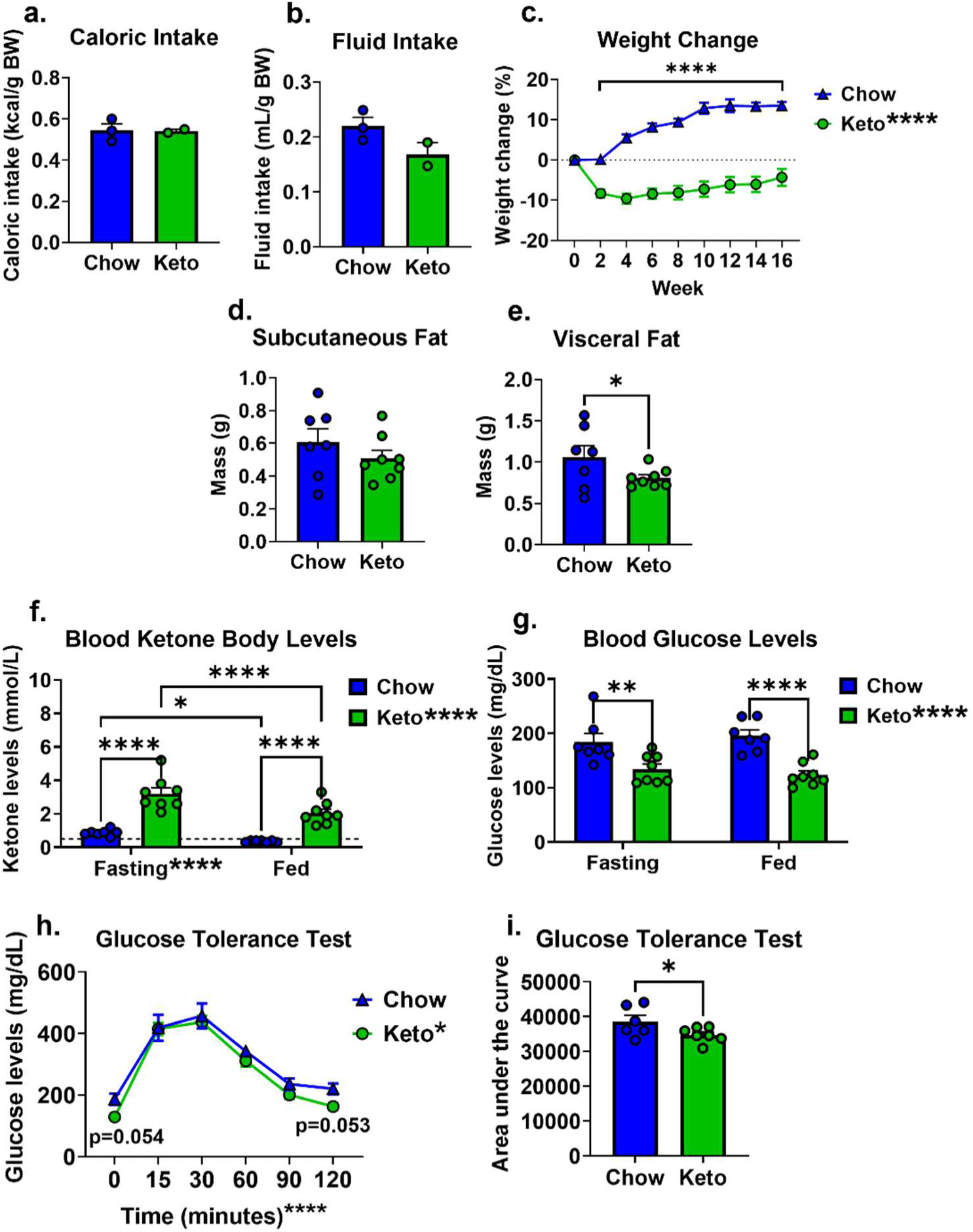
Ketogenic diet induced nutritional ketosis and had metabolic benefits in male Tg-SwDI mice. Caloric intake **(a)** and fluid intake **(b)** were measured biweekly, with data presented as an average value for each cage over the course of the diet intervention period. **(c)** Changes in body weight from baseline were measured throughout the diet intervention period. **(d)** Subcutaneous and **(e)** visceral fat were dissected and weighed at the time of euthanasia. **(f)** Ketone body and **(g)** glucose levels were measured in tail vein blood samples in both fasted and fed states. **(h)** Glucose tolerance testing was performed, with blood glucose levels measured at baseline in the fasted state (t=0), and at 15, 30, 60, 90, and 120 minutes after a glucose challenge injection. **(i)** Area under the curve was calculated for data collected during the glucose tolerance test as a measure of cumulative glucose exposure. *p<0.05, **p<0.01, ***p<0.001, ****p<0.0001.

#### Fluid intake

Fluid intake for each cage was approximated based on water bottle weights; these values were then normalized by average body weight of the mouse in that cage to calculate fluid intake in mL/g of body weight **(Figure 2b)**. An unpaired two-sample t-test found that mice fed a ketogenic diet tended to consume less fluid compared to mice on the standard chow diet; however, this was not significant, likely due to small sample sizes [t(3)=1.978, p=0.1423].

#### Body weight

Body weight change data are displayed as % change from baseline (prior to diet intervention) on a biweekly basis through the end of the experiment **(Figure 2c)**. A two-way repeated measures ANOVA found significant main effects of diet [F(1,104)=80.79, p<0.001; keto < chow] and time [F(8,104)=29.34, p<0.001; increased weight gain over time], as well as a diet x time interaction [F(8,104)=37.88, p<0.001]. Multiple pairwise comparisons revealed that chow-fed mice displayed significant weight gain from baseline in weeks 4 through 16 (p<0.0001 for all), while keto-fed mice displayed significant weight loss from baseline in weeks 2 through 16 (p<0.001).

#### Adiposity

Adiposity was assessed by measuring the mass of subcutaneous (**Figure 2d**) and visceral (**Figure 2e)** fat pads. While an unpaired two-sample t-test found no significant difference in subcutaneous fat between groups [t(13)=1.081, p=0.1497], another unpaired two-sample t-test found that mice fed a ketogenic diet had significantly less visceral fat compared to those on a standard chow diet [t(13)=1.810, p=0.0467].

#### Blood ketone body levels

Blood ketone body levels were measured in both the fasting and fed state to assess nutritional ketosis (**Figure 2f)**. One outlier was identified in the chow group for the measure of fed ketone body levels. A two-way repeated measures ANOVA found significant main effects of diet [F(1,13)=47.88, p<0.0001] and feeding state [F(1,13)=44.49, p<0.0001], as well as a diet x feeding state interaction [F(1,13)=6.648, p=0.0229]. Multiple pairwise comparisons found that ketone body levels were higher in the fasted versus the fed condition for both chow (p=0.0150) and keto (p<0.0001) mice, and that ketone body levels were higher in mice fed a ketogenic diet compared to mice fed a standard chow diet in both the fasting and fed conditions (p<0.0001 for both).

#### Blood glucose levels

Blood glucose levels were measured in both the fasting and fed state **(Figure 2g)**. A two-way repeated measures ANOVA found significant main effects of diet [F(1,13)=35.44, p<0.0001; keto < chow], while the main effect of feeding state [F(1,13)=0.0012, p=0.972] and the diet x feeding state interaction [F(1,13)=0.9059, p=0.3586]. Blood glucose levels were lower in mice fed a ketogenic diet compared to mice fed a standard chow diet in both the fasting (p=0.0034) and fed (p<0.0001) conditions.

#### Glucose tolerance test

Glucose tolerance testing was performed to assess diabetic status. Following measurement of fasting glucose levels, mice were injected with a bolus of glucose, with blood glucose levels measured at t = 15, 30, 60, and 120 min post-injection **(Figure 2h)**. One mouse was excluded from the chow group due to poor injection, and one outlier was identified in the ketogenic diet group. A two-way repeated measures ANOVA found significant main effects of diet [F(1,12)=5.467, p=0.0375; keto < chow] and time [F(5,60)=77.77, p<0.0001; expected rise and fall of glucose following challenge injection], while the diet x time interaction was not significant [F(5,60)=0.6205, p=0.6847]. Multiple pairwise comparisons revealed trends of attenuated blood glucose levels in mice fed a ketogenic diet compared to mice fed a standard chow diet at t=0 minutes (p=0.054) and t=120 minutes (p=0.053).

Additionally, area under the curve was calculated as a measure of cumulative glucose exposure during the glucose tolerance test **(Figure 2i)**. An unpaired two-sample t-test found that mice fed a ketogenic diet displayed significantly lower blood glucose levels compared to those on a standard chow diet [t(12)=2.007, p=0.0339].

### Behavior testing

#### Open field

The open field test was performed to assess general locomotor/exploratory behavior, as well as anxiety-like behavior **(Figure 3a-f)**. Unpaired two-sample t-tests performed separately for each outcome measure found that mice fed a ketogenic diet displayed significantly greater distance traveled [t(13)=2.819, p=0.0073, **Figure 3a**], time spent moving [t(13)=2.578 p=0.0115, **Figure 3b**], and velocity while moving [t(13)=2.352, p=0.0176, **Figure 3c**], while there were no group differences in time spent in the center of the arena [t(13)=0.4082, p=0.3449, **Figure 3d**], time spent rearing [t(13)=0.8735, p=0.1991, **Figure 3e**], nor time spent grooming [t(13)=0.8754, p=0.1986, **Figure 3f**].

**Figure 3.**
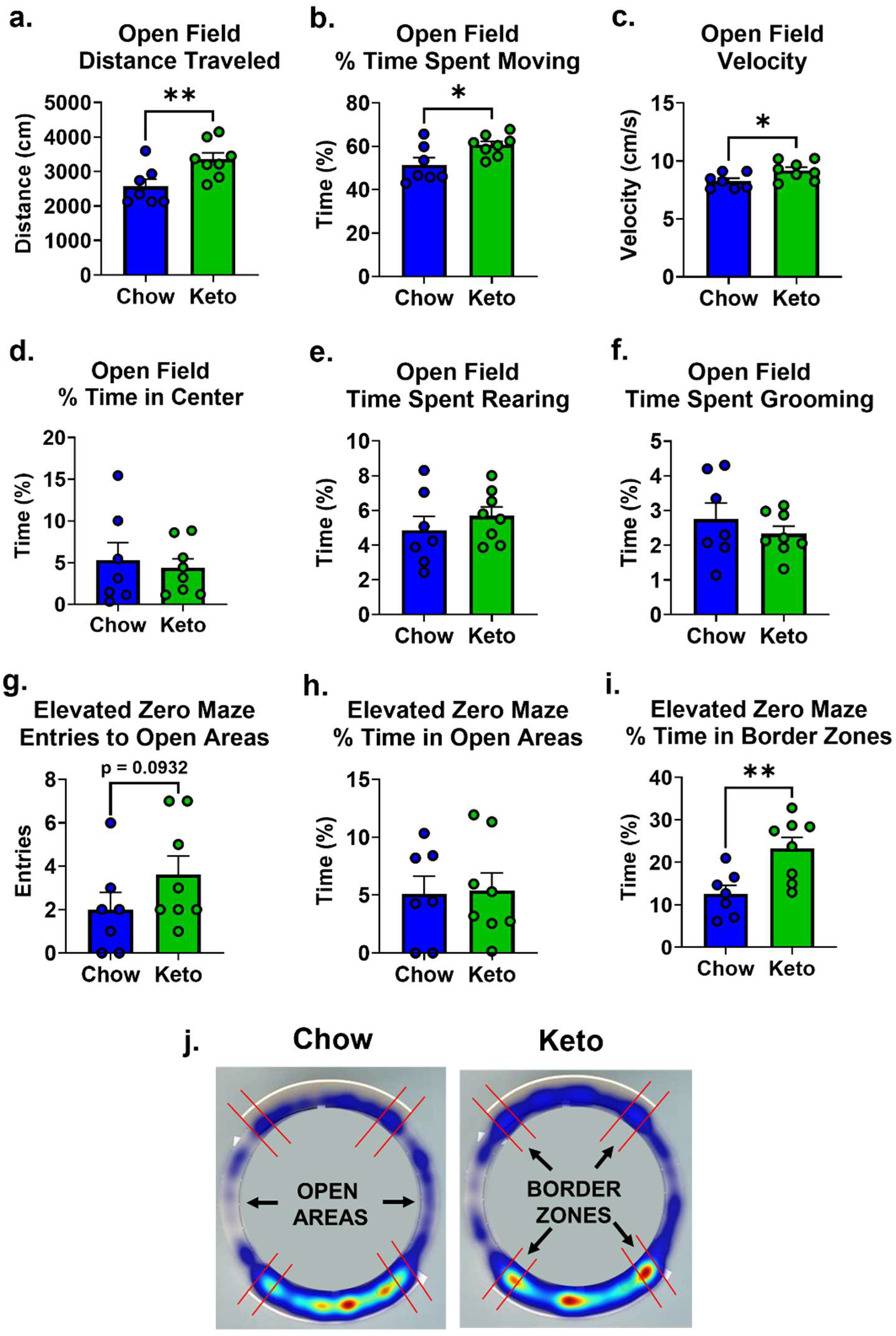
Ketogenic diet increased locomotor/exploratory activity without significantly affecting anxiety-like behavior in male Tg-SwDI mice. Ten-minute open field tests were conducted to assess general activity levels and anxiety-related behavior, including measures of **(a)** distance traveled, **(b)** time spent moving (%), **(c)** velocity, **(d)** center time (%), **(e)** rearing time (%), and **(f)** grooming time (%). Five-minute elevated zero maze tests were conducted to assess exploratory vs. anxiety-like behavior, including measures of **(g)** number of entries to open areas, **(h)** time spent in open areas (%), and **(i)** time spent in border zones (areas within the closed areas that immediately border the open areas). **(j)** A heatmap of average activity for each group during the elevated zero maze test. *p<0.05, **p<0.01.

#### Elevated zero maze

The elevated zero maze test was performed to assess anxiety-like behavior **(Figure 3g-i)**. Unpaired two-sample t-tests performed separately for each outcome measure revealed a trend that mice fed a ketogenic diet made more entries to the open areas compared to mice fed a standard chow diet [t(13)=1.395, p=0.0932, **Figure 3g**]; however, there were no group differences in time spent in the open areas [t(13)=0.1383, p=0.4460, **Figure 3h**].

Additionally, time spent in the border zones of the closed areas closest to the open areas was assessed, with an unpaired two-sample t-test revealing that mice fed a ketogenic diet spent more time in these border zones compared to mice on a standard chow diet [t(13)=3.185, p=0.0036, **Figure 3i**]. A heat map of activity in the elevated zero maze for the mean of each treatment group is shown in **Figure 3j**.

#### Novel object recognition test (NORT)

A novel object recognition test was performed to assess object recognition memory. There was no effect of diet on time spent exploring both objects (data not shown). Performance was quantified by the discrimination index **(Figure 4a)**. One outlier was identified in the ketogenic diet-fed group. One sample t-tests found that both chow [t(6)=5.969, p=0.0010] and keto [t(7)=7.211, p=0.0002] fed mice exhibited a significant preference for the novel object. An unpaired two-sample t-test found no difference in discrimination index between treatment groups [t(13)=0.5959, p=0.5245], suggesting that the ketogenic did not further enhance object recognition memory.

**Figure 4.**
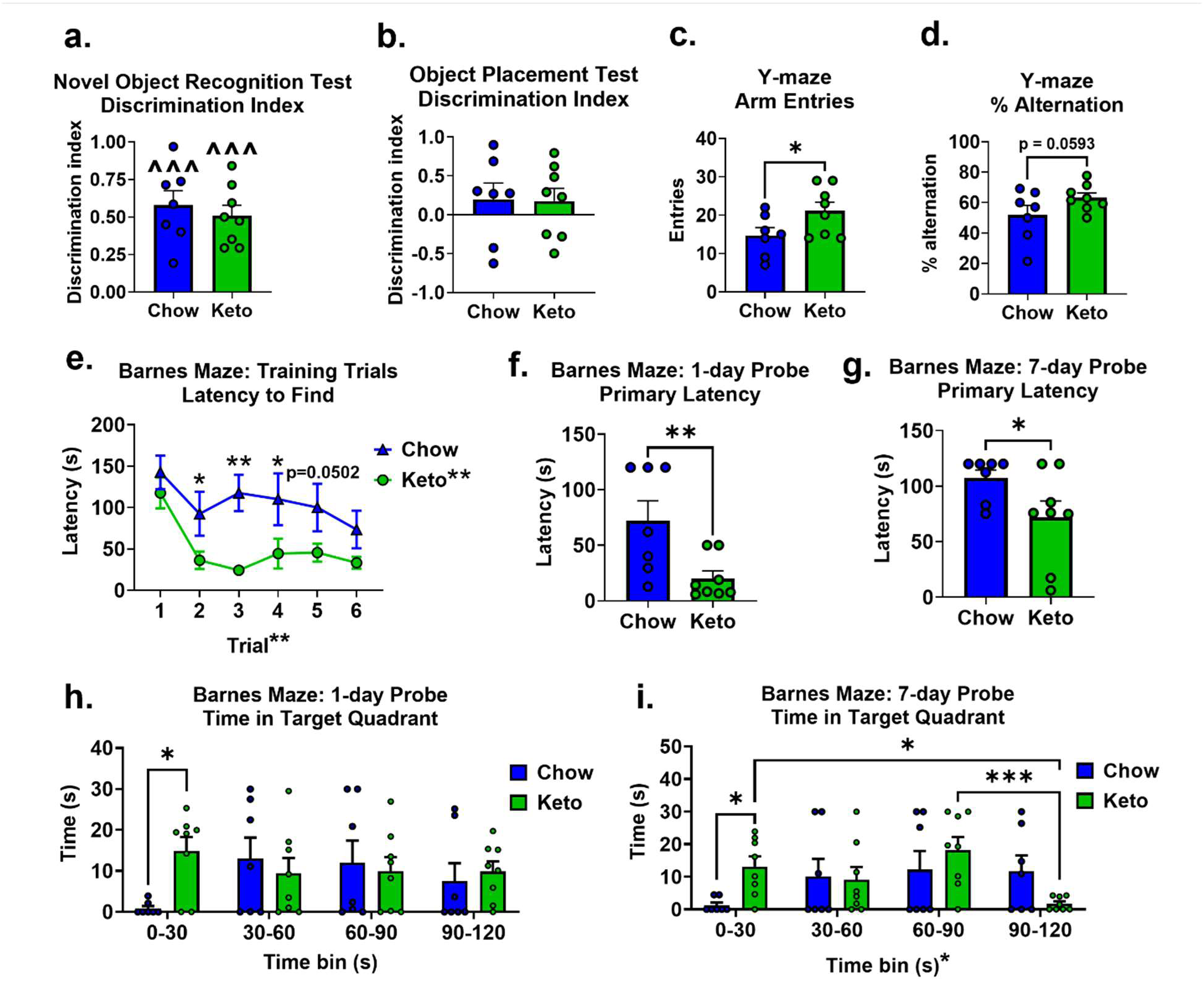
Ketogenic diet improved spatial learning and memory in male Tg-SwDI mice. **(a)** Novel object recognition testing was performed to assess object recognition memory, with training and testing trials separated by one hour. Time spent with the familiar and novels objects was measured during the five-minute testing trial, and discrimination indices were calculated as a measure of intact memory. **(b**) Object placement testing was performed to assess spatial memory, with training and testing trials separated by one hour. Time spent with the objects in the familiar and displaced locations was measured during the five-minute testing trial, and discrimination indices were calculated as a measure of intact memory. The y-maze test was performed to assess spatial working memory, with measures of **(c)** arm entries and **(d)** % alternation. The Barnes maze was conducted to assess spatial learning and memory. **(e)** Latency to find the target hole was measured over six training trials (two trials per day over three days) to assess spatial learning (maximum latency = 180 seconds). Primary latency, the latency to the first target hole visit, was measured to assess shorter- and longer-term memory during probe trails **(f)** one and **(g)** seven days after the last training trials (maximum latency = 120 seconds). Additionally, for the analysis of probe trials, the Barnes platform was divided into a center zone (30 cm diameter), with the remaining area split into 4 quadrants Time spent in the target quadrant was measured during the **(h)** 1-day and **(i)** 7-day probe trials. *p<0.05, **p<0.01, ***p<0.001. ^^^p<0.001 significantly greater than chance performance (hypothetical value of 0).

#### Object placement test (OPT)

An object placement test was performed to assess spatial memory. There was no effect of diet on time spent exploring both objects (data not shown). Performance was quantified by the discrimination index **(Figure 4b)**. One sample t-tests found that neither chow [t(6)=0.9500, p=0.3788] nor keto [t(7)=1.049, p=0.3289] fed mice exhibited a significant preference for the displaced object. An unpaired two-sample t-test found no difference in discrimination index between treatment groups [t(13)=0.0907, p=0.4645], suggesting that the ketogenic diet did not have any measurable benefit.

#### Y-maze

The Y-maze test was performed to assess spatial working memory **(Figure 4c-d)**. Unpaired two-sample t-tests performed separately for each outcome measure revealed that mice fed a ketogenic diet made more arm entries compared to mice fed a standard chow diet [t(13)=2.086, p=0.0286, **Figure 4c**], and that there was a trend of ketogenic diet increasing spatial working memory, as measure by % alternation [t(13)=1.671, p=0.0593, **Figure 4d**].

#### Barnes maze

The Barnes maze was performed to assess spatial learning and memory **(Figure 4e-i)**. Training trials took place over the course of 3 days with two trials per day, and latency to find the escape hole was measured **(Figure 4e)**. Across the six training trials, four outliers were identified amongst three different mice in the keto diet group. A two-way repeated measures ANOVA revealed main effects of diet [F(1,65)=17.39, p=0.0011; keto < chow] and time [F(5,65)=4.116, p=0.0026, decreased latency over trials], with no significant diet x time interaction [F(65)=0.7321, p=0.6020]. Multiple pairwise comparisons revealed that mice fed a ketogenic diet found the escape hole more quickly than mice fed a standard chow diet on days 2-5 (p<0.05 for all except day 5 where p=0.0573, trend).

Barnes maze probe trials were performed 1 and 7 days after the last training trial to assess spatial memory. Primary latency was calculated as the latency to the first visit to the previous target hole location for each probe trial. One outlier was identified within the 1-day probe trial for a mouse on the ketogenic diet. Unpaired two-sample t-tests were performed separately for each probe trial. Mice fed a ketogenic diet were faster to locate the previous hole location on both probe days 1 [t(13)=2.869, p=0.0066, **Figure 4f**] and 7 [t(13)=2.043, p=0.0310, **Figure 4g**].

Additionally, for each probe trial, the % time spent in the target quadrant, where the escape hole was previously located, was calculated, with each 2-minute probe divided into 30-second time bins to determine exploratory patterns during the trial. For the 1-day probe **(Figure 4h)**, a two-way repeated measures ANOVA found that neither the main effect of diet [F(1,13)=0.8308, p=0.3786] nor time [F(3,39)=0.4136, p=0.7442] were significant; however, there was a trend of a diet x time interaction [F(3,39)=2.362, p=0.0861]. Multiple pairwise comparisons found that during the first 30 seconds of the 1-day probe trial, mice fed a ketogenic diet spent significantly more time in the target quadrant compared to mice fed a standard chow diet (p=0.0119). For the 7-day probe **(Figure 4i)**, a two-way repeated measures ANOVA found that while the main effect of diet was not significant [F(1,13)=0.1633, p=0.6927], there was a significant main effect of time [F(3,39)=3.738, p=0.0188] and a diet x time interaction [F(3,39)=5.500, p=0.0030]. Multiple pairwise comparisons found that during the first 30 seconds of the 7-day probe trial, mice fed a ketogenic diet spent significantly more time in the target quadrant compared to mice fed a standard chow diet (p=0.0361).

### Neuropathological outcomes

#### Vascular density

Immunohistochemistry was performed for the assessment of vascular density, as measured by the % area covered by Collagen-IV immunolabeling **(Figure 5)**. Measures were taken in the cortex, thalamus, and subregions of the hippocampus (subiculum, dentate gyrus, CA1, and CA3). There was one outlier identified in the cortex and one in the CA3 region in the ketogenic diet group. Unpaired two-sample t-tests were performed separately for each brain region. While there was an increase in vascular density in the dentate gyrus following chronic administration of ketogenic diet [t(13)=2.582, p=0.0228 , **Figure 5d**], there was no effect of diet on vascular density in the cortex [t(13)=0.8198, p=0.4271, **Figure 5a**], thalamus [t(13)=0.8565, p=0.4072, **Figure 5b**], subiculum [t(13)=0.0333, p=0.9739, **Figure 5c**], CA1 [t(13)=1.287, p=0.2206, **Figure 5e**], nor CA3 [t(13)=1.447, p=0.1715, **Figure 5f**].

**Figure 5.**
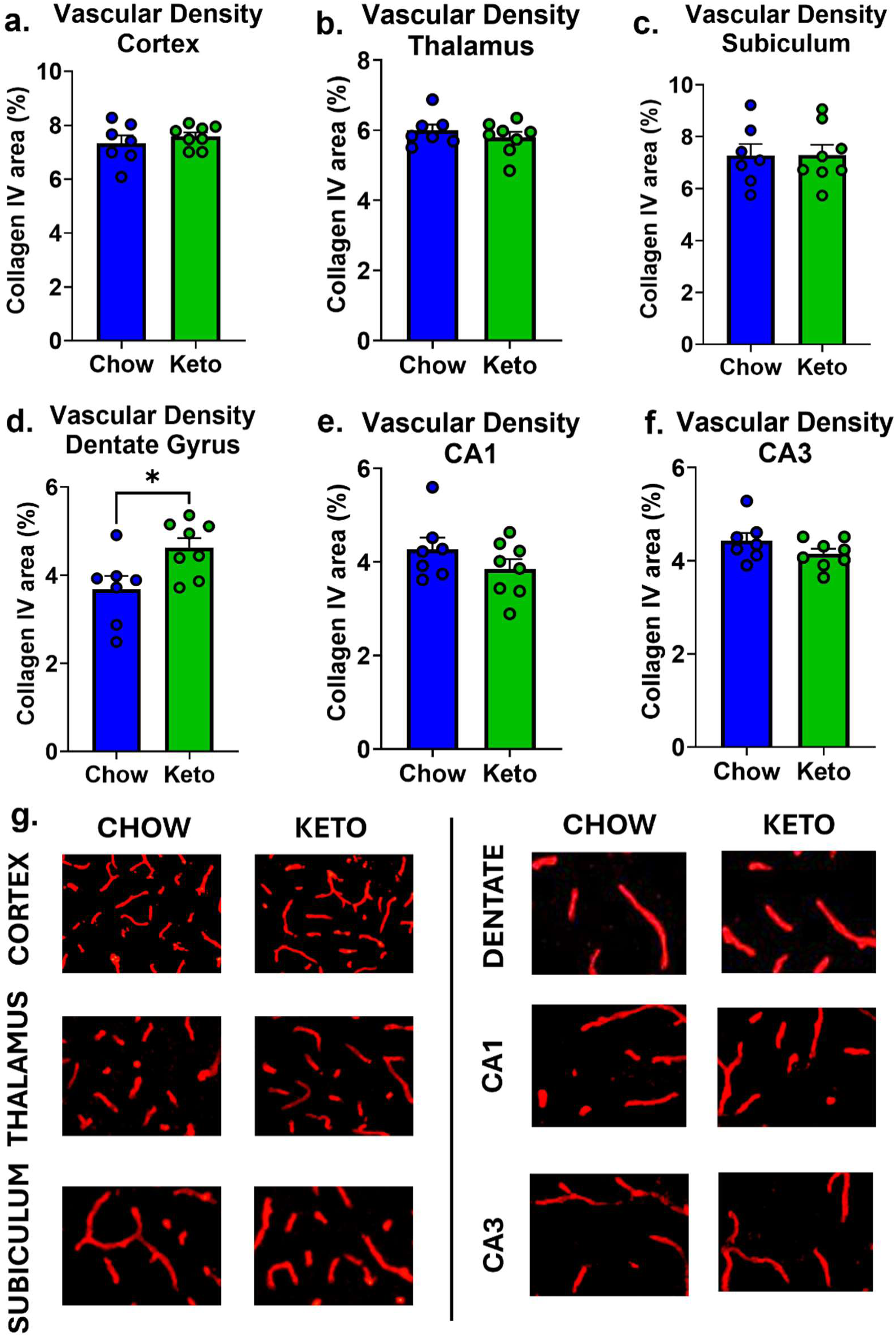
Ketogenic diet increased vascular density in the dentate gyrus only in male Tg-SwDI mice. Vascular density was assessed by quantifying the % area covered by Collagen IV immunolabeling in the **(a)** cortex, **(b)** thalamus, **(c)** subiculum, **(d)** dentate gyrus, **(e)** CA1, and **(f)** CA3. **(g)** Representative images of Collagen IV immunolabeling in each region of interest. *p<0.05.

#### Amyloid pathology

Staining with Thioflavin-S was performed for the localization and quantification of amyloid deposition in the cortex, thalamus, and subregions of the hippocampus (subiculum, dentate gyrus, CA1, and CA3) **(Figure 6)**. Assessment of overall amyloid pathology was measured by the total % area covered by Thioflavin-S staining in each region of interest. There was one dentate gyrus outlier in the chow-fed group. Unpaired two-sample t-tests were performed separately for each brain region. Ketogenic diet attenuated overall amyloid pathology in the cortex [t(13) = 2.090, p = 0.0284, **Figure 6a**] and dentate gyrus [t(13) = 4.189, p = 0.0006, **Figure 6d**], while there was a trend of decreased amyloid in the subiculum [t(13) = 1.494, p = 0.0795, **Figure 6c**]. There was no effect of diet on overall amyloid pathology in the thalamus [t(13)=1.120, p=0.1416, **Figure 6b**], CA1 [t(13)=0.5774, p=0.2868, **Figure 6e**], or CA3 [t(13)=1.171, p=0.1312, **Figure 6f**].

**Figure 6.**
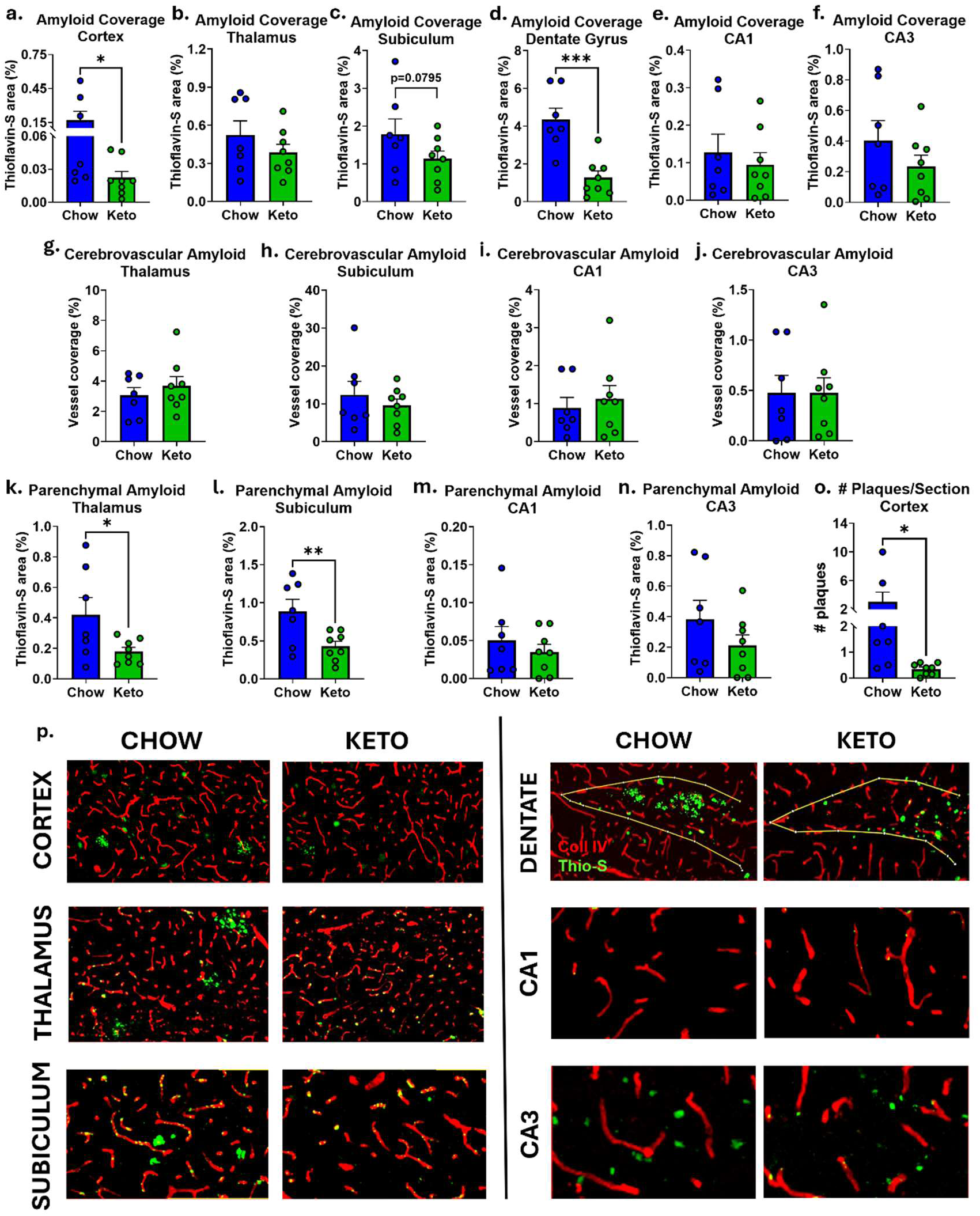
Ketogenic diet attenuated amyloid pathology, specifically plaque/parenchymal accumulation, in male Tg-SwDI mice. Thioflavin-S staining was performed to localize and quantify amyloid in the cortex, thalamus, and subregions of the hippocampus. **(a-f)** Overall amyloid pathology was quantified by measuring the % area covered by thioflavin-S staining in each region of interest. **(g-j)** Cerebrovascular amyloid, noted in the **(g)** thalamus, **(h)** subiculum, **(i)** CA1, and **(j)** CA3, was quantified by measuring the percentage of blood vessel area covered by amyloid. **(k-n)** Parenchymal amyloid deposition was quantified in brain regions exhibiting mixed plaque and CAA pathology. **(o)** Well-defined, thioflavin-S positive plaques were counted in the cortex. **(p)** Representative images of Collagen IV immunolabeling (red) and Thioflavin-S staining (green) in each region of interest. *p<0.05, **p<0.01, ***p<0.001.

While amyloid in the dentate gyrus and cortex was clearly deposited as plaque, additional measures were taken to determine the distribution of amyloid in the cerebral vasculature (CAA) and parenchyma in regions with potentially mixed distribution (thalamus, subiculum, CA1, CA3). Cerebral amyloid angiopathy was assessed by the % area of blood vessels covered by amyloid in a given brain region, as calculated by [(area of colocalized Thioflavin-S and Collagen IV)/(area of Collagen IV)]x100. Measures were taken in the thalamus and subregions of the hippocampus (subiculum, CA1, and CA3). One CA1 and one CA3 outlier was identified in the chow-fed group. Unpaired two-sample t-tests were performed separately for each brain region. There was no significant effect of ketogenic diet on CAA in the thalamus [t(13)=0.7688, p=0.2279, **Figure 6g**], subiculum [t(13)=0.7546, p=0.2320, **Figure 6h**], CA1 [t(13)=0.5232, p=0.30480 **Figure 6i**, nor CA3 [t(13)=0.0044, p=0.4983, **Figure 6j**].

Parenchymal (non-vascular) amyloid pathology was assessed by measuring the % area covered by Thio-flavin-S staining that was not colocalized with collagen IV immunolabeling (blood vessel marker). Measures were taken in the thalamus and subregions of the hippocampus (subiculum, CA1, and CA3) only, as amyloid pathology in the cortex and dentate were deemed to be plaque/parenchymal, with the overall amyloid measure sufficing for this purpose. One subiculum outlier and one CA1 outlier were identified in the ketogenic diet group. Unpaired two-sample t-tests were performed separately for each brain region. Ketogenic diet attenuated parenchymal (non-vascular) amyloid pathology in the thalamus [t(13)=2.188, p=0.0237, **Figure 6k**] and subiculum [t(13) = 2.781, p = 0.0078, **Figure 6l**] There was no effect of diet on parenchymal amyloid accumulation in the CA1 [t(13)=0.7452, p=0.2347, **Figure 6m**] nor CA3 [t(13)=1.253, p=0.1161, **Figure 6n**]. Additionally, plaque # was counted in the cortex, with one outlier identified in the ketogenic diet group. An unpaired two-sample t-test found that mice fed a ketogenic diet exhibited an attenuation in plaque # [t(13) = 2.156, p = 0.0252, **Figure 6o**].

#### Microgliosis

Immunolabeling for ionized calcium-binding adaptor molecule 1 (Iba1) was analyzed as a measure of microgliosis in the cortex, thalamus, and areas of the hippocampus (dentate gyrus, subiculum, CA1, CA3; **Figure 7**). One cortex outlier, one CA1 outlier and one CA3 outlier was identified in the chow-fed group. Unpaired two-sample t-tests were performed separately for each brain region. There was no effect of diet in the cortex [t(13)= 0.2818, p=0.7826, **Figure 7a**], thalamus [t(13)=1.434, p=0.1752, **Figure 7b**], subiculum [t(13)=0.5298, p=0.6052, **Figure 7c**], dentate gyrus [t(13)=0.3094, p=0.7619, **Figure 7d**], CA1 [t(13)=1.083, p=0.3453, **Figure 7e**], nor CA3 [t(13)=1.407, p=0.1828, **Figure 7f**].

**Figure 7.**
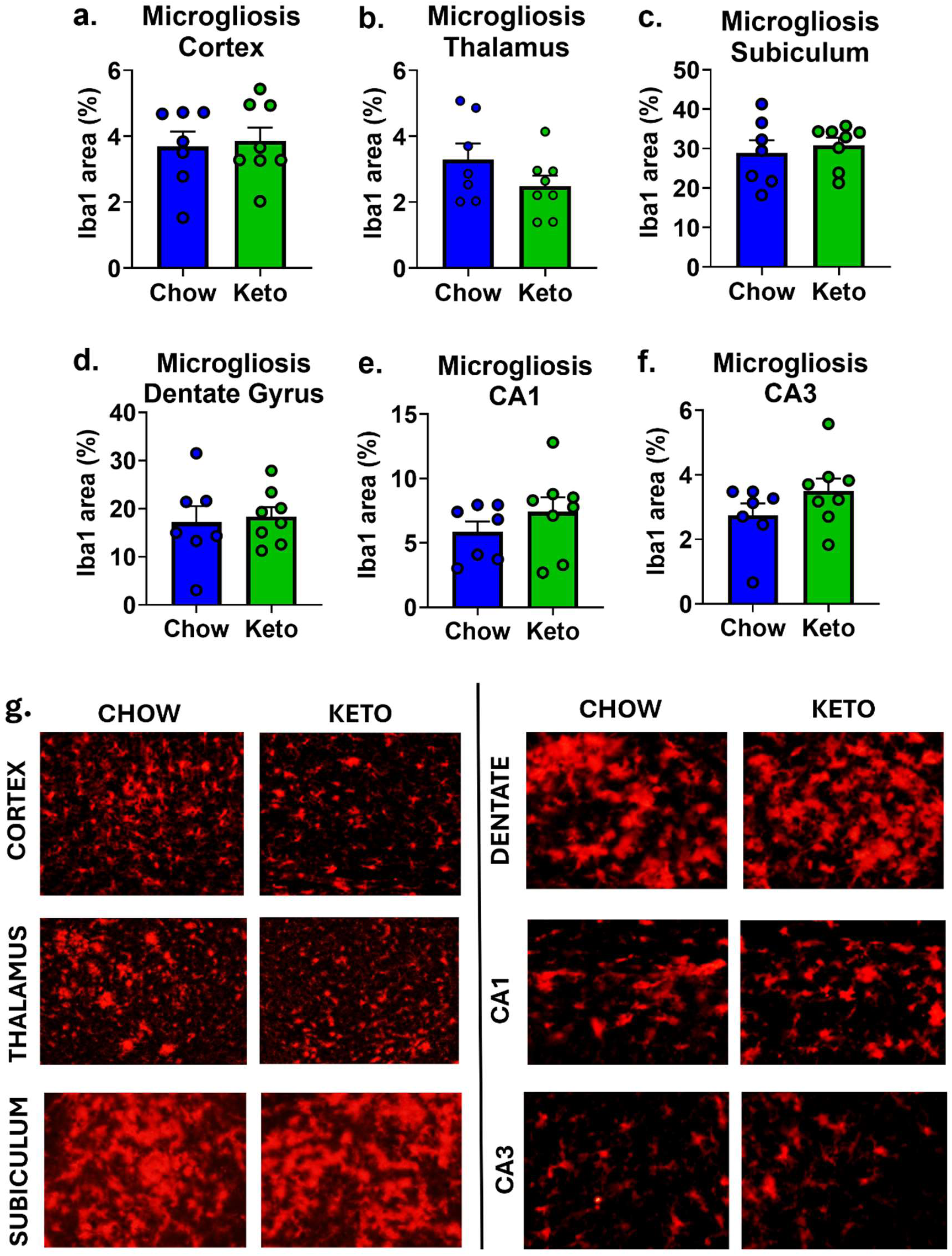
Ketogenic diet did not affect microgliosis in male Tg-SwDI mice. Microgliosis was assessed by quantifying the % area covered by ionized calcium binding adaptor molecule 1 (Iba1) immunolabeling in the **(a)** cortex, **(b)** thalamus, **(c)** subiculum, **(d)** dentate gyrus, **(e)** CA1, and **(f)** CA3. **(g)** Representative images of Iba1 immunolabeling in each region of interest.

#### Astrogliosis

Immunolabeling for glial fibrillary acidic protein (GFAP) was analyzed as a measure of astrogliosis in the cortex, thalamus, and areas of the hippocampus (dentate gyrus, subiculum, CA1, CA3). One mouse in the chow-fed group was identified as having outlier values in the thalamus and subiculum. Unpaired two-sample t-tests were performed separately for each brain region. In response to a ketogenic diet, there were trends of decreased GFAP immunolabeling in the cortex [t(13)=1.789, p=0.969, **Figure 8a**] and increased GFAP immunolabeling in the thalamus [t(13)=2.039, p=0.0623, **Figure 8b**], with no effect of diet in the subiculum [t(13)=1.562, p=0.1562, **Figure 8c**], dentate gyrus [t(13)=0.5668, p=0.5805, **Figure 8d**], CA1 [t(13)=1.347, p=0.2011, **Figure 8e**], nor CA3 [t(13)=0.4231, p=0.6791, **Figure 8f**].

**Figure 8.**
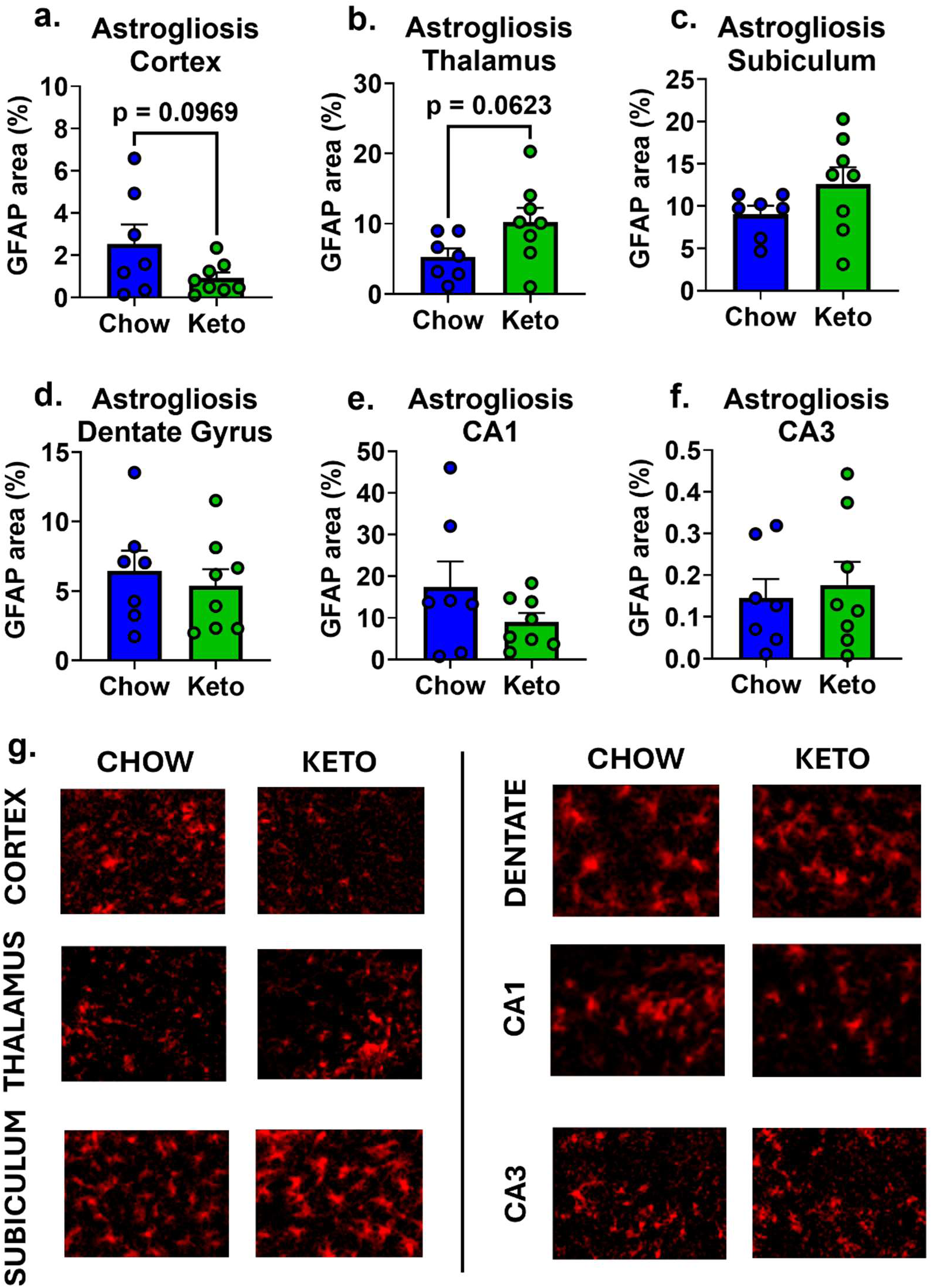
Ketogenic diet had negligible impact on astrogliosis in male Tg-SwDI mice. Astrogliosis was assessed by quantifying the % area covered by glial fibrillary acidic protein (GFAP) immunolabeling in the **(a)** cortex, **(b)** thalamus, **(c)** subiculum, **(d)** dentate gyrus, **(e)** CA1, and **(f)** CA3. **(g)** Representative images of GFAP immunolabeling in each region of interest.

#### Hippocampal Neurogenesis

Immunolabeling for doublecortin (DCX) was used to identify and count the number of immature neurons/neuroblasts in the dentate gyrus region of the hippocampus as a measure of neurogenesis. An unpaired two-sample t-test found that mice fed a ketogenic diet had a greater number of DCX+ cells compared to chow-fed mice [t(13)=3.073, p=0.0045], indicating an enhancement of neurogenesis **(Figure 9)**.

**Figure 9.**
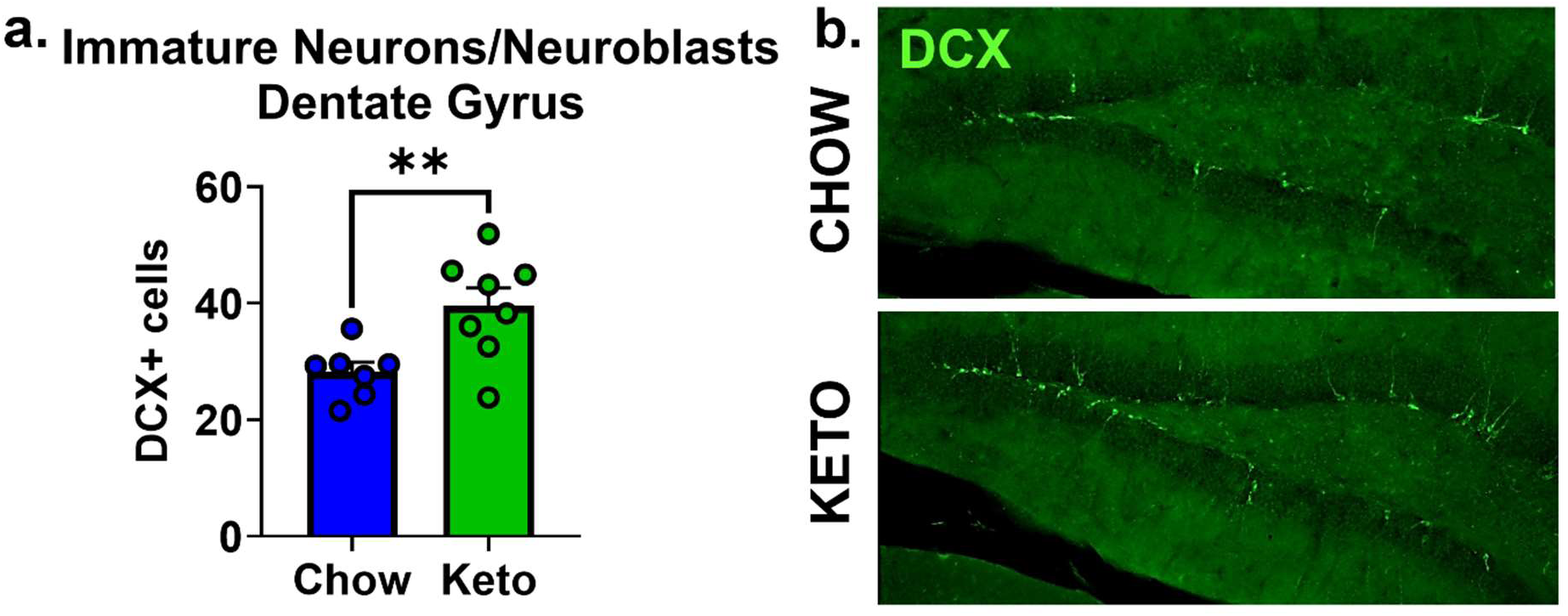
Ketogenic diet enhanced adult hippocampal neurogenesis in male Tg-SwDI mice. **(a)** Neurogenesis was assessed by counting the number of doublecortin (DCX)+ cells in the dentate gyrus of the hippocampus, as DCX-immunolabeling is indicative of immature neurons/neuroblasts. **(b)** Representative images of DCX immunolabeling in the dentate gyrus.

#### White matter

Immunolabeling for myelin basic protein (MBP) was used to characterize white matter integrity in the corpus callosum, cortex, and subregions of the hippocampus [dentate gyrus and stratum lacunosum-moleculare (SLM) layer]. There were no effects of diet on MBP intensity in any area measured, including the corpus callosum [t(13)=0.4245, p=0.6782, **Figure 10a**], cortex [t(13)=1.486, p=0.1611, **Figure 10b**], dentate gyrus of the hippocampus [t(13)=1.263, p=0.2287, **Figure 10c**], nor stratum lacunosum-moleculare (SLM) layer of the hippocampus [t(13)=0.5885, p=0.5663, **Figure 10d**].

**Figure 10.**
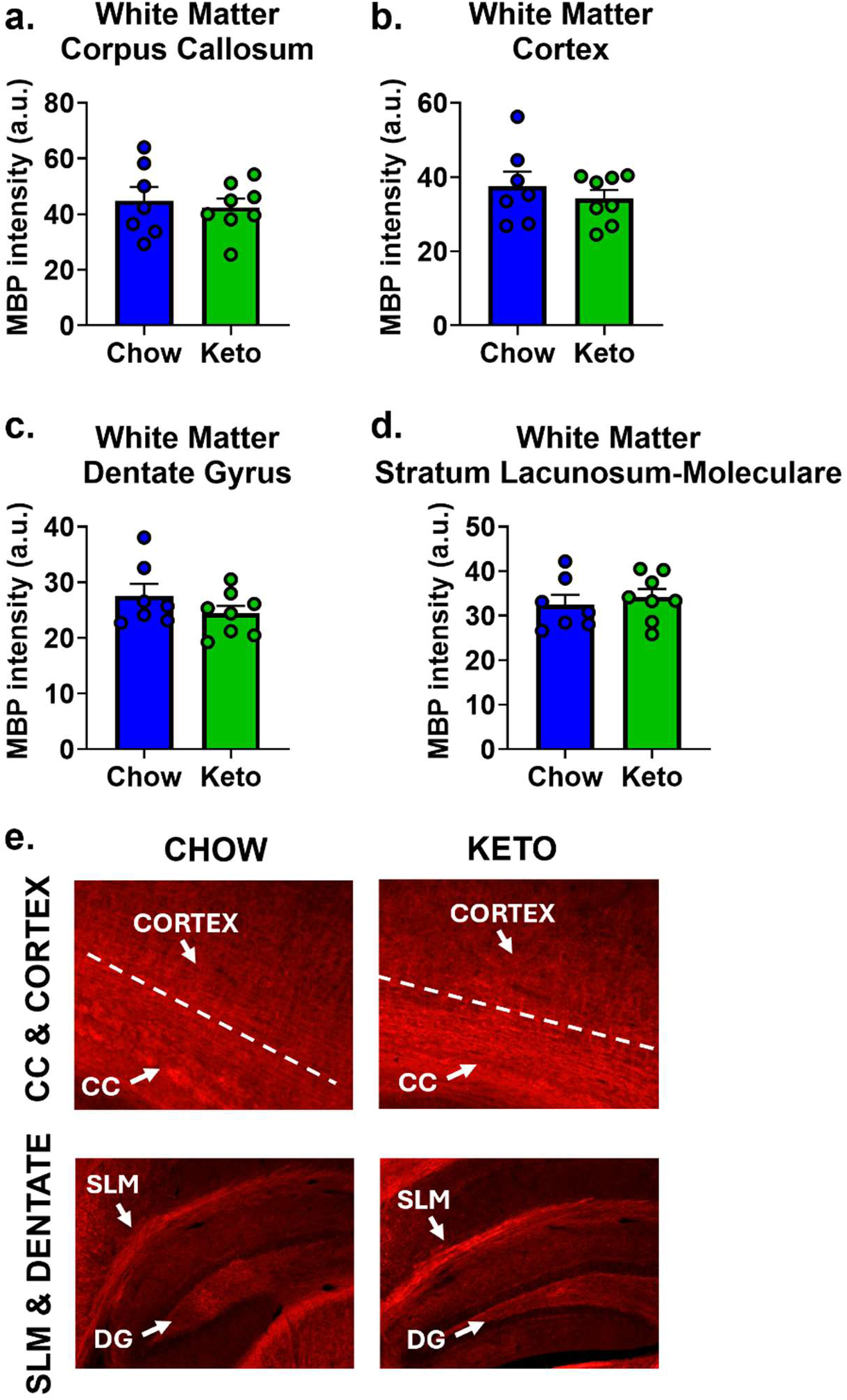
Ketogenic diet did not affect white matter integrity in male Tg-SwDI mice. White matter was assessed by measuring the intensity of MBP immunolabeling in the **(a)** corpus callosum (CC), **(b)** cortex, **(c)** dentate gyrus (DG), and **(d)** stratum lacunosum-moleculare (SLM). **(e)** Representative images of myelin basic protein (MBP) immunolabeling in each region of interest.

## DISCUSSION

Ketogenic dietary interventions have gained attention in recent years for their potential to combat neurodegenerative diseases, including Alzheimer’s disease (AD), as well as aid in the recovery from cerebrovascular events, such as stroke.18, 21, 23,24, 34 However, its utility for the prevention and/or treatment of dementia linked to cerebral amyloid angiopathy (CAA), a common form of cerebral small vessel disease that contributes to increased risk of dementia and stroke, had yet to be tested. Here, we fed male Tg-SwDI mice (a transgenic model exhibiting CAA) either standard chow or a ketogenic diet during early-to moderate-stage disease (∼3.5-7.5 months of age), followed by an assessment of metabolic, cognitive-behavioral, and neuropathological outcomes. We found that the ketogenic diet resulted in nutritional ketosis, improved several aspects of metabolic and cognitive-behavioral function, attenuated amyloid pathology, and enhanced adult hippocampal neurogenesis. These findings suggest that adherence to a ketogenic diet may provide a safe and effective means of combating dementia in patients with CAA.

The ketogenic diet is characterized by the consumption of a very low carbohydrate, moderate protein, and high fat diet, resulting in a state of “nutritional ketosis”. In the absence of sufficient carbohydrate intake, ketone bodies replace glucose as the primary source of energy. Nutritional ketosis can be evaluated by measuring blood ketone levels, typically defined by levels >0.5 mmol/L. Our ketogenic diet, with ∼3% carbohydrate content, achieved nutritional ketosis, with blood ketone levels exceeding this threshold, as well as levels observed in mice fed standard chow.

Despite its high fat content, the ketogenic diet has demonstrated numerous metabolic benefits, including weight loss, management of Type 2 diabetes and dyslipidemia, and attenuation of steatohepatitis.15, 16, 35 In line with these findings, we report that mice fed a ketogenic diet lost weight compared to baseline, while mice fed a standard chow diet gained weight. Adherence to the ketogenic diet also attenuated fasting and fed blood glucose levels while improving glucose tolerance. Furthermore, visceral fat was reduced, while subcutaneous fat was maintained by a ketogenic diet. Visceral fat, which accumulates around internal organs, is generally considered more harmful than subcutaneous fat, which is located beneath the skin. This difference in fat distribution is associated with varying inflammatory responses and metabolic consequences, with visceral fat generally being more harmful and linked to an increased risk of cardiometabolic and neurodegenerative diseases.36–38 Studies have shown that individuals with more visceral fat tend to have a higher risk of cognitive decline, reduced white matter integrity, and brain atrophy, particularly in regions critical for memory and cognition, such as the hippocampus.39–41 Visceral fat is metabolically active, secreting pro-inflammatory cytokines that increase systemic and neuro-inflammation, impair the blood-brain barrier, damage white matter, and contribute to insulin resistance, all of which are linked to cognitive decline and dementia.41–43 It is also more strongly associated with cardiovascular issues like atherosclerosis, which can reduce brain blood flow and increase the risk of dementia.44–46 In aging and dementia, weight loss and frailty are significant concerns, as they exacerbate physical and cognitive decline.47, 48 While the ketogenic diet led to mild early weight loss (<10%), weight began to rebound as the intervention continued. Moreover, subcutaneous fat levels were maintained, and activity levels increased, suggesting that the ketogenic diet may mitigate frailty and preserve physical function.

Beyond metabolic benefits, accumulating evidence highlights the ketogenic diet’s neuroprotective properties in mitigating neurodegenerative diseases and cerebrovascular events.49–53 In line with preclinical studies demonstrating the cognitive-preserving effects of ketogenic dietary interventions in mouse models of Alzheimer’s disease and cerebrovascular conditions,24, 54–56 we found that mice fed a ketogenic diet exhibited significant improvements across several cognitive-behavioral domains. Specifically, ketogenic diet-fed mice showed increased exploratory behavior (e.g., open field activity and y-maze arm entries), aligning with prior findings on the benefits of ketogenic supplements in other AD mouse models.57 Notably, we have previously shown that exploratory behavior is reduced in Tg-SwDI mice compared to wild-type controls,32, 58 suggesting that the observed increase likely reflects a restoration of healthy behavior rather than pathological hyperactivity. Anxiety-like behavior, assessed via the open field and elevated zero maze tests, was minimally affected by the ketogenic diet, with subtle increases in exploration of open areas potentially attributable to enhanced activity levels. In contrast, some previous studies have reported that ketogenic dietary interventions are anxiolytic in other AD mouse models.57, 59 In the current study, the ketogenic diet improved spatial working memory (y-maze spontaneous alternation), spatial learning, and short- and long-term spatial memory (Barnes maze). Performance on object-based tasks (novel object recognition and object placement tests) did not improve, likely due to task limitations (e.g., ceiling effects for novel object recognition and excessive difficulty for object placement). Previous studies have similarly reported enhancements in memory and learning tasks with ketogenic dietary interventions in rodent models of Alzheimer’s disease.52, 54, 55

These benefits to cognitive-behavioral function could be explained by the observed improvements in neuropathology and neuroplasticity. We found that the ketogenic diet reduced amyloid pathology, particularly plaque/parenchymal amyloid pathology in the cortex, thalamus, and sub-regions of the hippocampus, including the subiculum and dentate gyrus. This finding aligns with previous studies in rodent models, which have reported reductions in amyloid with ketogenic interventions, potentially contributing to cognitive-behavioral improvements by alleviating synaptic dysfunction and neuronal stress.55, 59, 60 There are several mechanisms by which a ketogenic diet could attenuate amyloid pathology,61 including neuroimmune mechanisms,62 though we did not observe significant changes in microglia and astrocyte immunolabeling. This diet also improves neuronal energy metabolism by providing ketone bodies as an alternative fuel, enhancing mitochondrial function, and reducing oxidative stress.52, 61, 63 The ketogenic diet may shift amyloid precursor protein (APP) processing toward a non-amyloidogenic pathway, which could contribute to an overall attenuation of Aβ production. Additionally, improved insulin sensitivity and restored glucose metabolism previously demonstrated to result from ketogenic interventions may enhance Aβ clearance via proteases like insulin-degrading enzyme.64 Of note, while non-vascular cerebral amyloid was reduced by the ketogenic diet, the level of vascular amyloid remained unchanged.

Although the ketogenic diet was initiated relatively early in the course of disease, vascular deposition may be less dynamic or reversible compared to parenchymal amyloid. Previously, we found that while exercise intervention from ∼4-12 months of age exerted cognitive-behavioral benefits and attenuated neuroinflammation, CAA levels were not reduced in mixed-sex Tg-SwDI mice.31 The persistence of vascular amyloid may be due to the distinct pathways and mechanisms involved in its deposition and clearance being less responsive to the ketogenic diet compared to mechanisms affecting parenchymal amyloid. Vascular amyloid, composed primarily of Aβ40, may accumulate earlier than parenchymal amyloid (predominantly Aβ42) and may be less responsive to metabolic shifts induced by the diet, though a previous study in AD mouse found both Aβ40 and Aβ42 to be reduced.60 Once vascular amyloid forms, even at early stages, it may act as a stable seed for further deposition, making it resistant to interventions.65 Early vascular dysfunction induced by CAA, including damage to smooth muscle and endothelial cells critical for Aβ clearance, may reduce the diet’s impact on vascular amyloid. Impaired perivascular drainage, a hallmark of CAA, limits the clearance of vascular amyloid, while enhanced glymphatic system activity demonstrated to result from ketogenic dietary interventions primarily affect parenchymal deposits.6, 66 Future studies should explore the contribution of these potential mechanisms and whether it is effective to combine ketogenic interventions with therapies that target vascular-specific mechanisms, such as enhancing perivascular drainage or addressing endothelial and smooth muscle cell dysfunction.

Mice fed the ketogenic diet showed increased adult hippocampal neurogenesis, evidenced by an increased number of immature neurons/neuroblasts in the dentate gyrus. Enhanced neurogenesis likely contributes to the observed improvements in spatial learning and memory, as it has been established that hippocampal neurogenesis is critical for cognitive function in this domain.67, 68 Although studies in healthy rodents have generally found no effect of ketogenic diet on neurogenesis,69, 70 these studies labeled different proteins (Ki67 or BrdU) for proliferating cells, compared to our labeling of immature neurons/neuroblasts using doublecortin (DCX). Other studies have demonstrated that ketogenic diet enhances neurogenesis in rodent models of neurological disease, suggesting that the diet may promote neurogenesis specifically in the context of injury, stress, or dysfunction and underscoring the diet’s potential to restore neuroplasticity. A ketogenic diet may increase neuroblasts and immature neurons in the dentate gyrus by creating a neuroprotective environment through multiple mechanisms. These include enhanced energy metabolism via ketone bodies, improved mitochondrial function, reduced oxidative stress and inflammation, modulation of epigenetic pathways, such as histone deacetylase (HDAC) inhibition, and upregulation of brain-derived neurotrophic factor (BDNF), which supports neurogenesis and synaptic plasticity.71–73

Interestingly, the ketogenic diet did not appear to confer measurable benefits for neuroinflammation, as indicated by immunofluorescence of astrocytes and microglia, nor for white matter integrity, as assessed by immunofluorescence of myelin basic protein. This contrasts with some studies that have reported reductions in neuroinflammation and improved white matter integrity following ketogenic interventions in other models of disease or injury.24, 34, 53, 54, 74–76 This may be due to variations in experimental models, disease stages, and/or dietary formulations.

This study has several limitations that should be addressed in future research. First, we employed a transgenic mouse model of CAA, which predominantly models hereditary disease, whereas most human CAA cases are sporadic. Additionally, the mouse model does not fully replicate all aspects of the human condition. Future studies should incorporate diverse animal models and eventually clinical trials to validate the generalizability of our findings. Second, our study included only male mice, leaving the potential influence of sex unexplored. There are reported sex differences in cognitive-behavioral performance and neuropathology in Tg-SwDI mice, with females being more adversely affected by the disease.58, 77, 78 We have previously shown striking sex differences in response to dietary interventions in other dementia mouse models.79–81 Evidence suggests that biological factors like age and sex differences can affect responses to ketogenic diets in both healthy and diseased states.54, 82–85 Follow-up studies should investigate the diet’s effects in both sexes to ensure broader applicability. Third, we initiated the ketogenic diet during early-stage disease. It remains unknown whether similar benefits would be observed in this model if the intervention were applied at later stages; however, a recent study found that ketogenic was effective in earlier but not later-stage disease in 5xFAD mice.54 Future research should evaluate the safety and efficacy of ketogenic diets at advanced disease stages of CAA. Finally, adherence to a ketogenic diet can be challenging, particularly for elderly individuals and those with dementia. To address this limitation, future studies should explore the potential of exogenous ketone supplementation, such as beta-hydroxybutyrate, as an alternative strategy. Preliminary evidence suggests that exogenous ketones may improve metabolic and cognitive outcomes in aging and Alzheimer’s disease, as well as reduce neuroinflammation, mitochondrial dysfunction, and reactive oxygen species, and support neuroplasticity.19–22 Further research is necessary to confirm these benefits and establish optimal dosing regimens.

## CONCLUSION

In conclusion, our findings demonstrate that a ketogenic diet can induce nutritional ketosis, improve metabolic and cognitive-behavioral functions, reduce amyloid pathology, and enhance neurogenesis in a transgenic mouse model of CAA. These results suggest that ketogenic dietary interventions hold promise as a non-pharmacological strategy for mitigating dementia in those affected with CAA. However, the limitations of this study highlight the need for additional research to optimize and translate these findings into clinical practice. By addressing these gaps, future studies can further elucidate the therapeutic potential of ketogenic diets and related interventions for neurodegenerative and cerebrovascular diseases.

## Author Contributions

Victoria Pulido-Correa (Conceptualization, Methodology, Data curation, Formal analysis, Funding acquisition, Investigation, Project administration, Visualization, Writing – original draft, Writing – review & editing); Ariana Hernandez (Investigation, Visualization, Writing – review & editing); Eleanor Wind (Methodology, Investigation, Writing – review & editing); Chana Vogel (Investigation, Writing – review & editing); YingYing Zhu (Investigation, Writing – review & editing); Shaina Binu (Investigation, Writing – review & editing); Mikayla Jeneske (Investigation, Writing – review & editing); Dylan Kuni (Investigation, Writing – review & editing); Vedika Chiduruppa (Investigation, Writing – review & editing); Bianca Echeverria (Investigation, Writing – review & editing); Lauren Rosenberg (Investigation, Writing – review & editing), Lisa Robison (Conceptualization, Methodology, Data curation, Formal analysis, Funding acquisition, Project administration, Supervision, Visualization, Writing – original draft, Writing – review & editing).

## Acknowledgments

The authors would like to thank Lauren White and Gabby Albensi for assistance with fluorescence imaging.

## Data Availability Statement

The authors confirm that the data supporting the findings of this study are available within the article. Raw data that support the findings of this study are available from the corresponding author upon reasonable request.

## REFERENCES

1. Arvanitakis Z, Leurgans SE, Wang Z, et al. Cerebral amyloid angiopathy pathology and cognitive domains in older persons. Ann Neurol 2011; 69: 320–327. 2011/03/10. DOI: 10.1002/ana.22112.

2. Wattendorff AR, Frangione B, Luyendijk W, et al. Hereditary cerebral haemorrhage with amyloidosis, Dutch type (HCHWA-D): clinicopathological studies. J Neurol Neurosurg Psychiatry 1995; 58: 699–705. 1995/06/01. DOI: 10.1136/jnnp.58.6.699.

3. Yamada M. Cerebral amyloid angiopathy: emerging concepts. J Stroke 2015; 17: 17–30. 2015/02/19. DOI: 10.5853/jos.2015.17.1.17.

4. Alakbarzade V, French JM, Howlett DR, et al. Cerebral amyloid angiopathy distribution in older people: A cautionary note. Alzheimer’s & Dementia: Translational Research & Clinical Interventions 2021; 7: e12145. DOI: 10.1002/trc2.12145.

5. Haussmann R, Homeyer P, Sauer C, et al. Comorbid cerebral amyloid angiopathy in dementia and prodromal stages-Prevalence and effects on cognition. Int J Geriatr Psychiatry 2023; 38: e6015. DOI: 10.1002/gps.6015.

6. Kim SH, Ahn JH, Yang H, et al. Cerebral amyloid angiopathy aggravates perivascular clearance impairment in an Alzheimer’s disease mouse model. Acta Neuropathol Commun 2020; 8: 181. 20201105. DOI: 10.1186/s40478-020-01042-0.

7. Greenberg SM, Bacskai BJ, Hernandez-Guillamon M, et al. Cerebral amyloid angiopathy and Alzheimer disease - one peptide, two pathways. Nat Rev Neurol 2020; 16: 30–42. 20191211. DOI: 10.1038/s41582-019-0281-2.

8. Salloway S, Chalkias S, Barkhof F, et al. Amyloid-Related Imaging Abnormalities in 2 Phase 3 Studies Evaluating Aducanumab in Patients With Early Alzheimer Disease. JAMA Neurol 2021 2021/11/23. DOI: 10.1001/jamaneurol.2021.4161.

9. Honig LS, Barakos J, Dhadda S, et al. ARIA in patients treated with lecanemab (BAN2401) in a phase 2 study in early Alzheimer’s disease. Alzheimers Dement (N Y) 2023; 9: e12377. 20230320. DOI: 10.1002/trc2.12377.

10. Cummings J, Apostolova L, Rabinovici GD, et al. Lecanemab: Appropriate Use Recommendations. J Prev Alzheimers Dis 2023; 10: 362–377. DOI: 10.14283/jpad.2023.30.

11. Sperling RA, Jack CR, Black SE, et al. Amyloid-related imaging abnormalities in amyloid-modifying therapeutic trials: recommendations from the Alzheimer’s Association Research Roundtable Workgroup. Alzheimer’s & dementia : the journal of the Alzheimer’s Association 2011; 7. DOI: 10.1016/j.jalz.2011.05.2351.

12. van Dyck CH, Sabbagh M and Cohen S. Lecanemab in Early Alzheimer’s Disease. Reply. N Engl J Med 2023; 388: 1631–1632. DOI: 10.1056/NEJMc2301380.

13. Greenberg SM, Cordonnier C, Schneider JA, et al. Off-label use of aducanumab for cerebral amyloid angiopathy. Lancet Neurol 2021; 20: 596–597. 2021/07/09. DOI: 10.1016/s1474-4422(21)00213-1.

14. Masood W, Annamaraju P, Khan Suheb MZ, et al. Ketogenic Diet. StatPearls. Treasure Island (FL): StatPearls Publishing Copyright © 2024, StatPearls Publishing LLC., 2024.

15. Luo W, Zhang J, Xu D, et al. Low carbohydrate ketogenic diets reduce cardiovascular risk factor levels in obese or overweight patients with T2DM: A meta-analysis of randomized controlled trials. Front Nutr 2022; 9: 1092031. 20221213. DOI: 10.3389/fnut.2022.1092031.

16. Zhou C, Wang M, Liang J, et al. Ketogenic Diet Benefits to Weight Loss, Glycemic Control, and Lipid Profiles in Overweight Patients with Type 2 Diabetes Mellitus: A Meta-Analysis of Randomized Controlled Trails. Int J Environ Res Public Health 2022; 19 20220822. DOI: 10.3390/ijerph191610429.

17. Chinna-Meyyappan A, Gomes FA, Koning E, et al. Effects of the ketogenic diet on cognition: a systematic review. Nutr Neurosci 2023; 26: 1258–1278. 20221110. DOI: 10.1080/1028415x.2022.2143609.

18. Tao Y, Leng SX and Zhang H. Ketogenic Diet: An Effective Treatment Approach for Neurodegenerative Diseases. Curr Neuropharmacol 2022; 20: 2303–2319. DOI: 10.2174/1570159x20666220830102628.

19. Makievskaya CI, Popkov VA, Andrianova NV, et al. Ketogenic Diet and Ketone Bodies against Ischemic Injury: Targets, Mechanisms, and Therapeutic Potential. International Journal of Molecular Sciences 2023; 24: 2576.

20. Saito ER, Warren CE, Hanegan CM, et al. A Novel Ketone-Supplemented Diet Improves Recognition Memory and Hippocampal Mitochondrial Efficiency in Healthy Adult Mice. Metabolites 2022; 12 20221025. DOI: 10.3390/metabo12111019.

21. Shippy DC, Wilhelm C, Viharkumar PA, et al. β-Hydroxybutyrate inhibits inflammasome activation to attenuate Alzheimer’s disease pathology. J Neuroinflammation 2020; 17: 280. 20200921. DOI: 10.1186/s12974-020-01948-5.

22. Youm YH, Nguyen KY, Grant RW, et al. The ketone metabolite β-hydroxybutyrate blocks NLRP3 inflammasome-mediated inflammatory disease. Nat Med 2015; 21: 263–269. 20150216. DOI: 10.1038/nm.3804.

23. Hersant H and Grossberg G. The Ketogenic Diet and Alzheimer’s Disease. J Nutr Health Aging 2022; 26: 606–614. DOI: 10.1007/s12603-022-1807-7.

24. Gibson CL, Murphy AN and Murphy SP. Stroke outcome in the ketogenic state--a systematic review of the animal data. J Neurochem 2012; 123 Suppl 2: 52–57. DOI: 10.1111/j.1471-4159.2012.07943.x.

25. Shaafi S, Mahmoudi J, Pashapour A, et al. Ketogenic Diet Provides Neuroprotective Effects against Ischemic Stroke Neuronal Damages. Adv Pharm Bull 2014; 4: 479–481. 20141231. DOI: 10.5681/apb.2014.071.

26. Percie du Sert N, Hurst V, Ahluwalia A, et al. The ARRIVE guidelines 2.0: Updated guidelines for reporting animal research. PLoS Biol 2020; 18: e3000410. 20200714. DOI: 10.1371/journal.pbio.3000410.

27. Davis J, Xu F, Deane R, et al. Early-onset and robust cerebral microvascular accumulation of amyloid beta-protein in transgenic mice expressing low levels of a vasculotropic Dutch/Iowa mutant form of amyloid beta-protein precursor. J Biol Chem 2004; 279: 20296–20306. 20040225. DOI: 10.1074/jbc.M312946200.

28. Miao J, Xu F, Davis J, et al. Cerebral microvascular amyloid beta protein deposition induces vascular degeneration and neuroinflammation in transgenic mice expressing human vasculotropic mutant amyloid beta precursor protein. Am J Pathol 2005; 167: 505–515. DOI: 10.1016/s0002-9440(10)62993-8.

29. Rosas-Hernandez H, Cuevas E, Raymick JB, et al. Impaired Amyloid Beta Clearance and Brain Microvascular Dysfunction are Present in the Tg-SwDI Mouse Model of Alzheimer’s Disease. Neuroscience 2020; 440: 48–55. 20200523. DOI: 10.1016/j.neuroscience.2020.05.024.

30. Xu F, Grande AM, Robinson JK, et al. Early-onset subicular microvascular amyloid and neuroinflammation correlate with behavioral deficits in vasculotropic mutant amyloid beta-protein precursor transgenic mice. Neuroscience 2007; 146: 98–107. 20070228. DOI: 10.1016/j.neuroscience.2007.01.043.

31. Robison LS, Popescu DL, Anderson ME, et al. Long-term voluntary wheel running does not alter vascular amyloid burden but reduces neuroinflammation in the Tg-SwDI mouse model of cerebral amyloid angiopathy. J Neuroinflammation 2019; 16: 144. 20190711. DOI: 10.1186/s12974-019-1534-0.

32. Robison LS, Francis N, Popescu DL, et al. Environmental Enrichment: Disentangling the Influence of Novelty, Social, and Physical Activity on Cerebral Amyloid Angiopathy in a Transgenic Mouse Model. Int J Mol Sci 2020; 21 20200128. DOI: 10.3390/ijms21030843.

33. Kraeuter AK, Guest PC and Sarnyai Z. The Y-Maze for Assessment of Spatial Working and Reference Memory in Mice. Methods Mol Biol 2019; 1916: 105–111. DOI: 10.1007/978-1-4939-8994-2_10.

34. Guo M, Wang X, Zhao Y, et al. Ketogenic Diet Improves Brain Ischemic Tolerance and Inhibits NLRP3 Inflammasome Activation by Preventing Drp1-Mediated Mitochondrial Fission and Endoplasmic Reticulum Stress. Front Mol Neurosci 2018; 11: 86. 20180320. DOI: 10.3389/fnmol.2018.00086.

35. Luukkonen PK, Dufour S, Lyu K, et al. Effect of a ketogenic diet on hepatic steatosis and hepatic mitochondrial metabolism in nonalcoholic fatty liver disease. Proc Natl Acad Sci U S A 2020; 117: 7347–7354. 20200316. DOI: 10.1073/pnas.1922344117.

36. Zhang S, Huang Y, Li J, et al. Increased visceral fat area to skeletal muscle mass ratio is positively associated with the risk of cardiometabolic diseases in a Chinese natural population: A cross-sectional study. Diabetes Metab Res Rev 2023; 39: e3597. 20221202. DOI: 10.1002/dmrr.3597.

37. Dolatshahi M, Commean PK, Rahmani F, et al. Alzheimer Disease Pathology and Neurodegeneration in Midlife Obesity: A Pilot Study. Aging Dis 2024; 15: 1843–1854. 20240801. DOI: 10.14336/ad.2023.0707.

38. Huang X, Wang YJ and Xiang Y. Bidirectional communication between brain and visceral white adipose tissue: Its potential impact on Alzheimer’s disease. EBioMedicine 2022; 84: 104263. 20220919. DOI: 10.1016/j.ebiom.2022.104263.

39. Raji CA, Meysami S, Hashemi S, et al. Visceral and Subcutaneous Abdominal Fat Predict Brain Volume Loss at Midlife in 10,001 Individuals. Aging Dis 2024; 15: 1831–1842. 20240801. DOI: 10.14336/ad.2023.0820.

40. Ozato N, Saitou S, Yamaguchi T, et al. Association between Visceral Fat and Brain Structural Changes or Cognitive Function. Brain Sci 2021; 11 20210804. DOI: 10.3390/brainsci11081036.

41. O’Brien PD, Hinder LM, Callaghan BC, et al. Neurological consequences of obesity. Lancet Neurol 2017; 16: 465–477. DOI: 10.1016/s1474-4422(17)30084-4.

42. Lampe L, Zhang R, Beyer F, et al. Visceral obesity relates to deep white matter hyperintensities via inflammation. Ann Neurol 2019; 85: 194–203. DOI: 10.1002/ana.25396.

43. Xu H, Barnes GT, Yang Q, et al. Chronic inflammation in fat plays a crucial role in the development of obesity-related insulin resistance. J Clin Invest 2003; 112: 1821–1830. DOI: 10.1172/jci19451.

44. Laurin D, Masaki KH, White LR, et al. Ankle-to-brachial index and dementia: the Honolulu-Asia Aging Study. Circulation 2007; 116: 2269–2274. 20071022. DOI: 10.1161/circulationaha.106.686477.

45. Xie B, Shi X, Xing Y, et al. Association between atherosclerosis and Alzheimer’s disease: A systematic review and meta-analysis. Brain Behav 2020; 10: e01601. 20200311. DOI: 10.1002/brb3.1601.

46. Neeland IJ, Ross R, Després JP, et al. Visceral and ectopic fat, atherosclerosis, and cardiometabolic disease: a position statement. Lancet Diabetes Endocrinol 2019; 7: 715–725. 20190710. DOI: 10.1016/s2213-8587(19)30084-1.

47. Halil M, Cemal Kizilarslanoglu M, Emin Kuyumcu M, et al. Cognitive aspects of frailty: mechanisms behind the link between frailty and cognitive impairment. J Nutr Health Aging 2015; 19: 276–283. DOI: 10.1007/s12603-014-0535-z.

48. Franx BAA, Arnoldussen IAC, Kiliaan AJ, et al. Weight Loss in Patients with Dementia: Considering the Potential Impact of Pharmacotherapy. Drugs Aging 2017; 34: 425–436. DOI: 10.1007/s40266-017-0462-x.

49. Rong L, Peng Y, Shen Q, et al. Effects of ketogenic diet on cognitive function of patients with Alzheimer’s disease: a systematic review and meta-analysis. J Nutr Health Aging 2024; 28: 100306. 20240628. DOI: 10.1016/j.jnha.2024.100306.

50. Wu L, Chu L, Pang Y, et al. Effects of dietary supplements, foods, and dietary patterns in Parkinson’s disease: meta-analysis and systematic review of randomized and crossover studies. Eur J Clin Nutr 2024; 78: 365–375. 20240220. DOI: 10.1038/s41430-024-01411-1.

51. Al-Kuraishy HM, Jabir MS, Albuhadily AK, et al. Role of ketogenic diet in neurodegenerative diseases focusing on Alzheimer diseases: The guardian angle. Ageing Res Rev 2024; 95: 102233. 20240214. DOI: 10.1016/j.arr.2024.102233.

52. Oliveira TPD, Morais ALB, Dos Reis PLB, et al. A Potential Role for the Ketogenic Diet in Alzheimer’s Disease Treatment: Exploring Pre-Clinical and Clinical Evidence. Metabolites 2023; 14 20231229. DOI: 10.3390/metabo14010025.

53. Jang J, Kim SR, Lee JE, et al. Molecular Mechanisms of Neuroprotection by Ketone Bodies and Ketogenic Diet in Cerebral Ischemia and Neurodegenerative Diseases. Int J Mol Sci 2023; 25 20231221. DOI: 10.3390/ijms25010124.

54. Xu Y, Jiang C, Wu J, et al. Ketogenic diet ameliorates cognitive impairment and neuroinflammation in a mouse model of Alzheimer’s disease. CNS Neurosci Ther 2022; 28: 580–592. 20211210. DOI: 10.1111/cns.13779.

55. Qin Y, Bai D, Tang M, et al. Ketogenic diet alleviates brain iron deposition and cognitive dysfunction via Nrf2-mediated ferroptosis pathway in APP/PS1 mouse. Brain Res 2023; 1812: 148404. 20230508. DOI: 10.1016/j.brainres.2023.148404.

56. Beckett TL, Studzinski CM, Keller JN, et al. A ketogenic diet improves motor performance but does not affect β-amyloid levels in a mouse model of Alzheimer’s disease. Brain Res 2013; 1505: 61–67. 20130212. DOI: 10.1016/j.brainres.2013.01.046.

57. Pawlosky RJ, Kashiwaya Y, King MT, et al. A Dietary Ketone Ester Normalizes Abnormal Behavior in a Mouse Model of Alzheimer’s Disease. Int J Mol Sci 2020; 21 20200204. DOI: 10.3390/ijms21031044.

58. Setti SE, Flanigan T, Hanig J, et al. Assessment of sex-related neuropathology and cognitive deficits in the Tg-SwDI mouse model of Alzheimer’s disease. Behav Brain Res 2022; 428: 113882. 20220406. DOI: 10.1016/j.bbr.2022.113882.

59. Kashiwaya Y, Bergman C, Lee JH, et al. A ketone ester diet exhibits anxiolytic and cognition-sparing properties, and lessens amyloid and tau pathologies in a mouse model of Alzheimer’s disease. Neurobiol Aging 2013; 34: 1530–1539. 20121229. DOI: 10.1016/j.neurobiolaging.2012.11.023.

60. Van der Auwera I, Wera S, Van Leuven F, et al. A ketogenic diet reduces amyloid beta 40 and 42 in a mouse model of Alzheimer’s disease. Nutr Metab (Lond) 2005; 2: 28. 20051017. DOI: 10.1186/1743-7075-2-28.

61. Taylor MK, Sullivan DK, Keller JE, et al. Potential for Ketotherapies as Amyloid-Regulating Treatment in Individuals at Risk for Alzheimer’s Disease. Front Neurosci 2022; 16: 899612. 20220616. DOI: 10.3389/fnins.2022.899612.

62. Jorfi M, Maaser-Hecker A and Tanzi RE. The neuroimmune axis of Alzheimer’s disease. Genome Med 2023; 15: 6. 20230126. DOI: 10.1186/s13073-023-01155-w.

63. Pinto A, Bonucci A, Maggi E, et al. Anti-Oxidant and Anti-Inflammatory Activity of Ketogenic Diet: New Perspectives for Neuroprotection in Alzheimer’s Disease. Antioxidants 2018; 7: 63.

64. Farris W, Mansourian S, Chang Y, et al. Insulin-degrading enzyme regulates the levels of insulin, amyloid β-protein, and the β-amyloid precursor protein intracellular domain in vivo. Proceedings of the National Academy of Sciences 2003; 100: 4162–4167.

65. Xu F, Fu Z, Dass S, et al. Cerebral vascular amyloid seeds drive amyloid β-protein fibril assembly with a distinct anti-parallel structure. Nat Commun 2016; 7: 13527. 20161121. DOI: 10.1038/ncomms13527.

66. Tan X, Li X, Li R, et al. β-hydroxybutyrate alleviates neurological deficits by restoring glymphatic and inflammation after subarachnoid hemorrhage in mice. Exp Neurol 2024; 378: 114819. 20240517. DOI: 10.1016/j.expneurol.2024.114819.

67. Lieberwirth C, Pan Y, Liu Y, et al. Hippocampal adult neurogenesis: Its regulation and potential role in spatial learning and memory. Brain Res 2016; 1644: 127–140. 20160510. DOI: 10.1016/j.brainres.2016.05.015.

68. Frechou MA, Martin SS, McDermott KD, et al. Adult neurogenesis improves spatial information encoding in the mouse hippocampus. Nat Commun 2024; 15: 6410. 20240730. DOI: 10.1038/s41467-024-50699-x.

69. Strandberg J, Kondziella D, Thorlin T, et al. Ketogenic diet does not disturb neurogenesis in the dentate gyrus in rats. Neuroreport 2008; 19: 1235–1237. DOI: 10.1097/WNR.0b013e32830a7109.

70. Irfannuddin I, Sarahdeaz SFP, Murti K, et al. The effect of ketogenic diets on neurogenesis and apoptosis in the dentate gyrus of the male rat hippocampus. J Physiol Sci 2021; 71: 3. 20210119. DOI: 10.1186/s12576-020-00786-7.

71. Pinto A, Bonucci A, Maggi E, et al. Anti-Oxidant and Anti-Inflammatory Activity of Ketogenic Diet: New Perspectives for Neuroprotection in Alzheimer’s Disease. Antioxidants (Basel) 2018; 7 20180428. DOI: 10.3390/antiox7050063.

72. Kim SW, Marosi K and Mattson M. Ketone beta-hydroxybutyrate up-regulates BDNF expression through NF-κB as an adaptive response against ROS, which may improve neuronal bioenergetics and enhance neuroprotection (P3.090). Neurology 2017; 88: P3.090. DOI: doi:10.1212/WNL.88.16_supplement.P3.090.

73. Wang X, Wu X, Liu Q, et al. Ketogenic metabolism inhibits histone deacetylase (HDAC) and reduces oxidative stress after spinal cord injury in rats. Neuroscience 2017; 366: 36–43.

74. Jiang J, Pan H, Shen F, et al. Ketogenic diet alleviates cognitive dysfunction and neuroinflammation in APP/PS1 mice via the Nrf2/HO-1 and NF-κB signaling pathways. Neural Regen Res 2023; 18: 2767–2772. DOI: 10.4103/1673-5374.373715.

75. Mu J, Wang T, Li M, et al. Ketogenic diet protects myelin and axons in diffuse axonal injury. Nutr Neurosci 2022; 25: 1534–1547. 20210123. DOI: 10.1080/1028415x.2021.1875300.

76. Stumpf SK, Berghoff SA, Trevisiol A, et al. Ketogenic diet ameliorates axonal defects and promotes myelination in Pelizaeus-Merzbacher disease. Acta Neuropathol 2019; 138: 147–161. 20190327. DOI: 10.1007/s00401-019-01985-2.

77. Maniskas ME, Mack AF, Morales-Scheihing D, et al. Sex differences in a murine model of Cerebral Amyloid Angiopathy. Brain Behav Immun Health 2021; 14: 100260. 20210417. DOI: 10.1016/j.bbih.2021.100260.

78. Finger C, Lee J and Manwani B. Sex Differences in Cerebral Amyloid Angiopathy: The Role of Monocytes/Macrophages (S2.008). Neurology 2022; 98: 3443. DOI: doi:10.1212/WNL.98.18_supplement.3443.

79. Robison LS, Gannon OJ, Thomas MA, et al. Role of sex and high-fat diet in metabolic and hypothalamic disturbances in the 3xTg-AD mouse model of Alzheimer’s disease. J Neuroinflammation 2020; 17: 285. 20200929. DOI: 10.1186/s12974-020-01956-5.

80. Gannon OJ, Robison LS, Salinero AE, et al. High-fat diet exacerbates cognitive decline in mouse models of Alzheimer’s disease and mixed dementia in a sex-dependent manner. J Neuroinflammation 2022; 19: 110. 20220514. DOI: 10.1186/s12974-022-02466-2.

81. Salinero AE, Robison LS, Gannon OJ, et al. Sex-specific effects of high-fat diet on cognitive impairment in a mouse model of VCID. Faseb j 2020; 34: 15108–15122. 20200916. DOI: 10.1096/fj.202000085R.

82. Cochran J, Taufalele PV, Lin KD, et al. Sex Differences in the Response of C57BL/6 Mice to Ketogenic Diets. Diabetes 2018; 67. DOI: 10.2337/db18-1884-P.

83. Eap B, Nomura M, Panda O, et al. Ketone body metabolism declines with age in mice in a sex-dependent manner. bioRxiv 2022: 2022.2010.2005.511032. DOI: 10.1101/2022.10.05.511032.

84. Kumar M, Bhatt B, Gusain C, et al. Sex-specific effects of ketogenic diet on anxiety-like behavior and neuroimmune response in C57Bl/6J mice. J Nutr Biochem 2024; 127: 109591. 20240202. DOI: 10.1016/j.jnutbio.2024.109591.

85. Modica LCM, Flores-Felix K, Casachahua LJD, et al. Impact of ketogenic diet and ketone diester supplementation on body weight, blood glucose, and ketones in Sprague Dawley rats fed over two weeks. Food Chem (Oxf) 2021; 3: 100029. 20210610. DOI: 10.1016/j.fochms.2021.100029.

